# Systematically quantifying morphological features reveals constraints on organoid phenotypes

**DOI:** 10.1101/2021.01.08.425947

**Authors:** Lauren E. Beck, Jasmine Lee, Christopher Coté, Margaret C. Dunagin, Ilya Lukonin, Nikkita Salla, Marcello K. Chang, Alex J. Hughes, Joseph D. Mornin, Zev J. Gartner, Prisca Liberali, Arjun Raj

**Affiliations:** Department of Bioengineering, School of Engineering and Applied Sciences, University of Pennsylvania, Philadelphia, PA, USA; Department of Genetics, Perelman School of Medicine, University of Pennsylvania, Philadelphia, PA, USA; Friedrich Miescher Institute for Biomedical Research (FMI), Basel, CH; University of Basel, Basel, CH; Department of Cell and Developmental Biology, Perelman School of Medicine, University of Pennsylvania, Philadelphia, PA, USA; Independent Researcher, Berkeley, CA, USA; Department of Pharmaceutical Chemistry, University of California, San Francisco, San Francisco, CA, USA; Center for Cellular Construction, University of California, San Francisco, San Francisco, CA, USA; Chan Zuckerberg Biohub, San Francisco, CA, USA

## Abstract

Organoids recapitulate complex 3D organ structures and represent a unique opportunity to probe the principles of self-organization. While we can alter an organoid’s morphology by manipulating the culture conditions, the morphology of an organoid often resembles that of its original organ, suggesting that organoid morphologies are governed by a set of tissue-specific constraints. Here, we establish a framework to identify constraints on an organoid’s morphological features by quantifying them from microscopy images of organoids exposed to a range of perturbations. We apply this framework to Madin-Darby Canine Kidney cysts and show that they obey a number of constraints taking the form of scaling relationships or caps on certain parameters. For example, we found that the number, but not size, of cells increases with increasing cyst size. We also find that these constraints vary with cyst age and can be altered by varying the culture conditions. We observed similar sets of constraints in intestinal organoids. This quantitative framework for identifying constraints on organoid morphologies may inform future efforts to engineer organoids.

## Introduction

Organoids are 3D structures that grow entirely *in vitro* from single or small groups of cells that mimic organ anatomy. Organoids have the potential to transform both personalized and regenerative medicine, since thousands of organoids can be grown under controlled conditions *in vitro* from small amounts of donor tissue. It is clear that organoids can form intricate biological structures, and these structures have an overall structure that resembles the associated organ. Yet, at the same time, there is often enormous variability between individual organoids (Garreta et al. 2020; Koo et al. 2019; Phipson et al. 2019; Kim, Koo, and Knoblich 2020; Volpato et al. 2018), and changing the organoid culture protocol can similarly lead to large changes (Yin et al. 2014; Sidhaye and Knoblich 2020; Gjorevski et al. 2016). Thus, the question remains as to what constraints organoids obey to give rise to the aspects of their morphology that are immutable, and what aspects of their morphology are either variable or tunable. Categorizing organoid features in this way may help reveal the design principles underlying organoid development.

One way to formalize the concept of constraints is via the dimensionality of morphospace, which is the set of morphologies an organism or model system can have. If one were to measure all possible features of an organoid’s morphology (e.g. size, number of nuclei, together comprising the axes of an organoid’s morphospace) across a large enough number of organoids, it could be that a large number of these features would strongly covary and thus could be explained by a single variable. For example, if size and number of nuclei were to be strongly correlated, then the dimensionality would effectively be 1 instead of 2, and the relationship between these variables would constitute a constraint on organoid morphology. On the other hand, if variables show a lack of correlation, then that would suggest independent axes of variability, indicating an additional degree of freedom in organoid morphospace. Examples of such dimension reduction have been demonstrated in both *C. elegans* and Snapdragon flowers, four dimensions capture over 90% of the variance in morphologies (Stephens et al. 2008; Cui et al. 2010). However, such analyses have not been performed in organoids yet. Recent work has quantitatively described brain organoid morphologies (Albanese et al. 2020) and uncovered genetic interactions governing intestinal organoid morphologies (Lukonin et al. 2020), but the constraints on organoid morphologies have not been characterized.

One potential reason that there are few quantitative analyses of organoid morphologies is that previous studies have been limited to a small set of two-dimensional features, such as organoid area and nuclear intensity, that fail to fully capture many characteristic aspects of the organoid’s shape (Kassis et al. 2019; Gracz et al. 2015). A major challenge is that quantifying morphological features such as the number of cells, cell shapes, etc., often requires microscopy images to be annotated to outline each individual cell or nucleus. While algorithms for automatic segmentation for images of large three-dimensional structures are improving (Piccinini et al. 2020), in many instances, segmentation must still be done manually to ensure sufficient accuracy. Such issues are compounded in organoids with many cell types and complex three-dimensional structures that are difficult to quantitatively align and compare to each other. Simpler “model” organoid systems might serve as a proving ground to test concepts about morphospaces.

How might we characterize the constraints on organoid morphologies? Our approach was to use variation in organoid morphology-both naturally occurring and variation induced by external stimuli-to sample the organoid morphospace. As a proof of concept, we developed this approach in spherical cysts grown from Madin-Darby Canine Kidney (MDCK) cells. We then quantified morphological features (cyst size, number of cells, eccentricity, etc) and the relationships between them (how does the number of cells scale with cyst size), thus revealing the constraints on MDCK cyst morphologies. To overcome the challenge of systematically quantifying morphological features, we combined algorithms for generating candidate annotations with software that allowed for quick manual correction. We found that MDCK cyst morphologies all fell along a small set of dimensions. These dimensions encoded a number of constraints; for instance, larger cysts had increased number but not size of constituent cells. We also found that some of these constraints on MDCK cyst morphologies vary with age and can be perturbed through drugs and growth factors. We used quantitative data on enteroid size, cell number, and cell type composition to reveal similar constraints in that more complex organoid system. Our results demonstrate a general strategy for determining the ways in which organoid morphologies are either constrained or free to vary.

## Results

### MOCK cyst morphologies span a limited number of dimensions

To quantify constraints on cyst morphologies, we designed an experimental and analytical workflow for culturing cysts, performing 3D imaging, annotating structures of interest, and measuring morphological features (Fig. 1A). We chose to apply this approach to MDCK cysts because of their relative simplicity and because they are amenable to high magnification 3D imaging. MDCK cells are an immortalized epithelial line that grow in adherent culture on 2D substrates, but form hollow 3D cysts when cultured in 3D matrices such as collagen or Matrigel (Supp. Fig. 1). A MDCK cyst grows from a single cell and is composed of an outer layer of polarized cells surrounding one to many lumens. The combination of their simplicity with the existence of a number of structural features to quantify make MDCK cysts an ideal system for establishing a framework for quantifying constraints on organoid morphologies.

**Figure 1:**
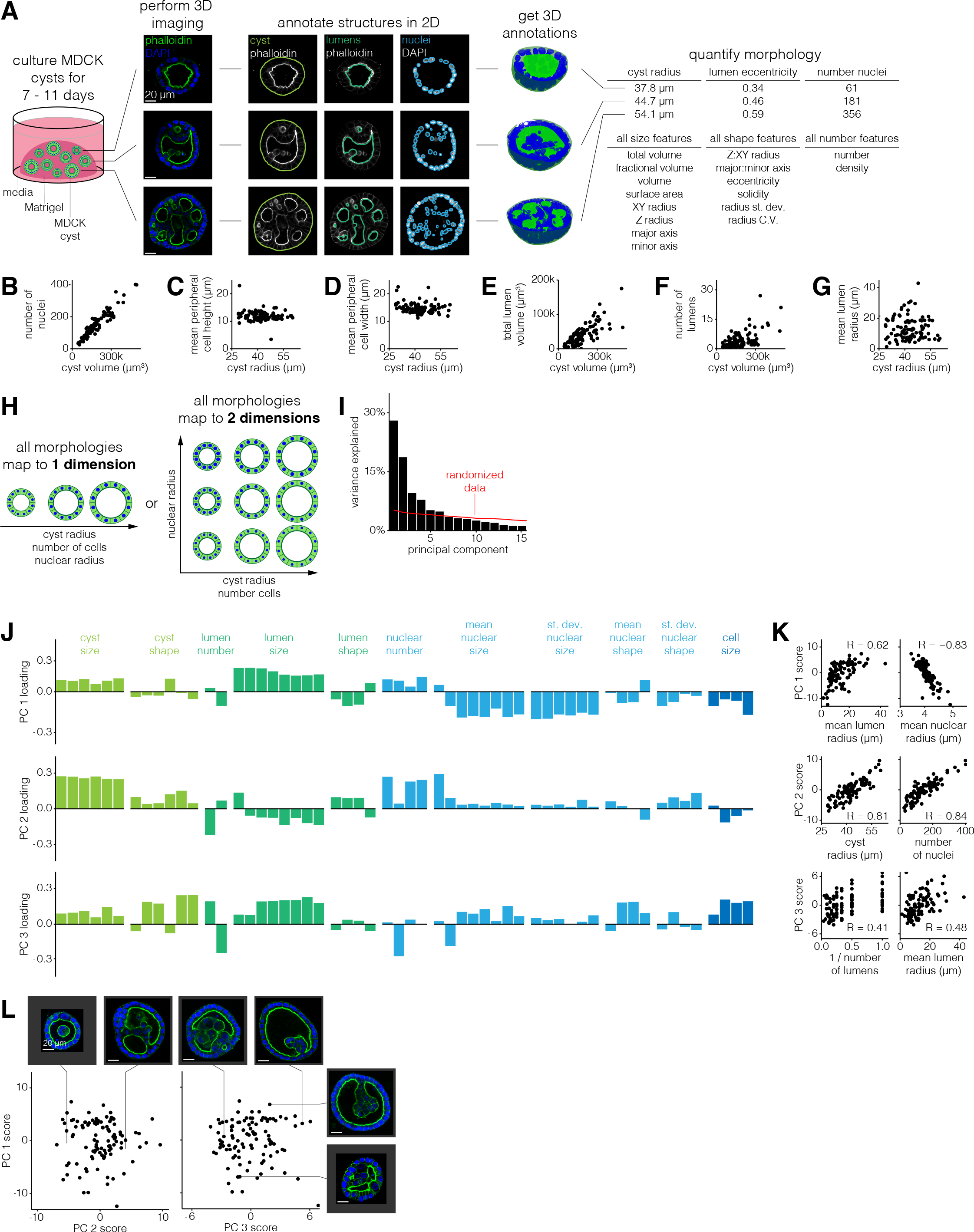
MOCK cyst morphologies span a limited number of dimensions. **A.** Schematic of experiments to quantify MOCK cyst morphological features. Briefly, we culture cysts for a variable number of days, perform 3O imaging of nuclei and cell boundaries for at least half of the cyst, annotate the boundaries of the cyst and each nucleus and lumen, and measure morphological features on the 3O annotations. **B-G.** Comparison of two morphological features for a time-window of 7-11 day old MOCK cysts. **H.** Example schematic of MOCK cyst morphologies that are captured by 1 dimension or 2 dimensions. **I.** Variance explained by each principal component. The red line indicates how much variance is explained when the data is randomized before PCA (see methods for details). **J.** Loading of each feature on principal components one through three. Each feature is color-coded by what structure (cyst, lumen, nucleus, or cell) it describes. **K.** Correlation between principal component score and raw morphological features for features which highly contribute to that principal component. **L.** Principal component scores for the first three principal components. Each pair of example cysts were chosen because they have high vs low score for one principal component, and similar scores for the other two principal components.

(We evaluated other organoid systems, such as the gut organoids, for our analysis, but found that the complexity of their morphologies presented a much larger challenge. For instance, gut organoids have complex bud structures that one would need to align to each other for quantitative comparison. The comparative simplicity of MDCK cysts that are at least nominally spherical made our analysis feasible as a proof of principle, with MDCK cysts serving as a model for more complex organoids.) In order to quantify 3D measurements of morphological features for hundreds of MDCK cysts, we established a pipeline for semi-automatically annotating cyst structures (nuclei, lumens, and cyst boundary). To identify the lumen and cyst boundaries, we developed a custom analysis pipeline that generated candidate annotations and accepted manual corrections to the annotations as needed (Supp. Fig. 2). We used cellpose (Stringer et al. 2020) to identify the boundaries of each nucleus and a custom analysis pipeline to manually correct the 3D annotations (Supp. Fig. 3). Due to depth of field limitations, in many cases we could not image the full depth of the cyst. We thus made sure to image at least the bottom half of the cyst, from which we computationally determined the middle point of each cyst and measured features on only the bottom half of each cyst to ensure a fair comparison between all cysts. We then measured morphological features of size, shape, and number on the nuclei, lumen, and cyst annotations anticipating that these may be the features that vary amongst cysts (Table 1). We performed a variety of comparisons between morphological features in order to verify that our measurements were consistent with basic geometric constraints. For example, because the cysts always appeared spherical, we confirmed that the cyst volume scaled with the cube of the cyst radius (Supp. Fig. 4).

We also confirmed that the total lumen and total nuclear volume was always less than that of the cyst volume, and we visually inspected cysts with high and low feature metrics to confirm that the quantified differences reflected differences in the images (Supp. Fig. 5-6).

We then wanted to find relationships between features that could potentially reflect biological constraints. For example, did the number of cells scale with the size of the cyst, as is typically the case in mammalian systems (Hafen and Stocker 2003; Savage et al. 2007)? Or, did larger cysts have the same number of cells as smaller cysts, but with larger component cells? We used the number of nuclei as a proxy for the number of cells and found that larger cysts had proportionally more nuclei (Fig. 1B). Because cells peripheral to the lumen(s) had different morphology than those internal to the lumens, we wondered whether their number scaled differently with cyst volume. We found that the number of peripheral nuclei scaled sublinearly with cyst volume (Supp. Fig. 7). Surprisingly, the number of internal cells scaled superlinearly with cyst volume, thus ensuring that the total number of cells scaled linearly with cyst volume. Given that the number of cells scaled with the cyst volume, we predicted that cell size should be independent of cyst size. We found that despite increases in cyst volume the peripheral cell height and width are fairly constant at -9-13 µm and -12-18 µm, respectively (Fig. 1C-D). Together, we called this set of constraints the constant-cell-density constraint.

We also wondered how lumens, both in number and size, scaled with increasing cyst size. For example, if a cyst is larger must it also have larger lumens? One alternative is that there is maximum lumen size, and larger cysts then have multiple lumens of the same size as smaller cysts. We found that the total lumen volume increased with increasing cyst volume, but that this could be achieved through one large lumen or many smaller lumens (Fig. 1E-G). However, we did notice that there was seemingly a maximum number of lumens per cyst that increased linearly with cyst volume. We called this constraint the “lumen number cap”, and its existence suggests that there may be a minimum lumen size (Supp. Fig. 8).

Given that MDCK cysts obey a number of constraints, we then wondered whether these constraints are coupled. In other words, might there be a single dimension (or a few dimensions), each of which may comprise several correlated features, along which all MDCK cyst morphologies fall (Fig. 1H)? To identify dimensions in the space of MDCK cyst morphologies we performed principal component analysis (PCA) on the set of 77 cysts and their 66 morphological features. In order to apply PCA to our data, we needed to supply a single value for each feature for each cyst. For all features describing nuclei we used both the mean and standard deviation across all nuclei within the cyst, e.g. mean nuclear volume and standard deviation of nuclear volume. For features describing lumens we used the mean across all lumens in the cyst, e.g. mean lumen volume. We didn’t include other higher order statistics like standard deviation because it was impossible to do so for the many cysts that had only one lumen. We found that the first three principal components respectively explain 28%, 19%, and 10% of the variation in MDCK cyst morphologies (Fig. 1I). We then wondered whether the principal components reflected any of the constraints we had previously identified. We found that the first principal component represented lumen size and inversely nuclear size, reflecting the fact that as lumens get larger, nuclei get smaller (Fig. 1J-L, Supp. Fig. 9). Note that this trend is consistent with our earlier finding of relatively constant cell size because cell size and nuclear size are different properties. The second principal component represents cyst size and number of nuclei, reflecting that increased cyst size was associated with increased number of nuclei, a relationship we previously identified as the constant-cell-density constraint. The third principal component represented the trade-off between lumen size and the number of lumens, reflecting that, for a given cyst size, in order to have more lumens, the individual lumens must be smaller (rather than there is a maximum lumen size which is independent of the number of lumens). The third principal component also represented a trade-off between nuclear size and the density of nuclei, reflecting that, for a given cyst size, in order to have more nuclei the nuclei must be smaller (Supp. Fig. 10). (Consistent with PC1, we also found that nuclear size anti-correlated with lumen size.) Beyond those three principal components, the remaining components accounted for less variation than components calculated from randomized data, suggesting that those PCs likely do not reflect substantial variation in the data. In addition to performing conventional PCA on our data, we also used a sparse PCA method (Benjamin Erichson et al. 2018), which can aid with interpretation because it tries to reduce small contributions to principal components to zero when possible. We found that the principal components from such an analysis (after discarding the first principal component, which is most likely technical and batch variability) recapitulated similar contributions to the axes of variation revealed by conventional PCA. This analysis also had a more straightforward interpretation owing to the principal components having just a few primary contributing features (Supp. Fig. 11). Thus, despite quantifying a large number of features, MDCK cyst morphologies can thus be represented by a limited number of dimensions.

### Constraints on MOCK cyst morphologies vary with age

MDCK cysts grow continuously over the course of weeks, from a single cell into large cysts. To determine whether or not cyst age affected the constraints on MDCK cyst morphologies, we compared the quantitative relationship between various features for cysts of different ages. We partitioned our data based on cyst age, ranging from 3-17 days of growth. Using the same imaging and feature quantification described above, we saw that, as expected, old cysts were larger in volume on average (Fig. 2A-B). We also noticed that, for a given age, there was a spread in cyst sizes. This variation in size enabled us to compare constraints in cyst size across different age categories (Fig. 2C). We used cysts cultured for 9 days as a reference point for younger and older cysts to evaluate how constraints on MDCK cyst morphologies changed with cyst age.

**Figure 2:**
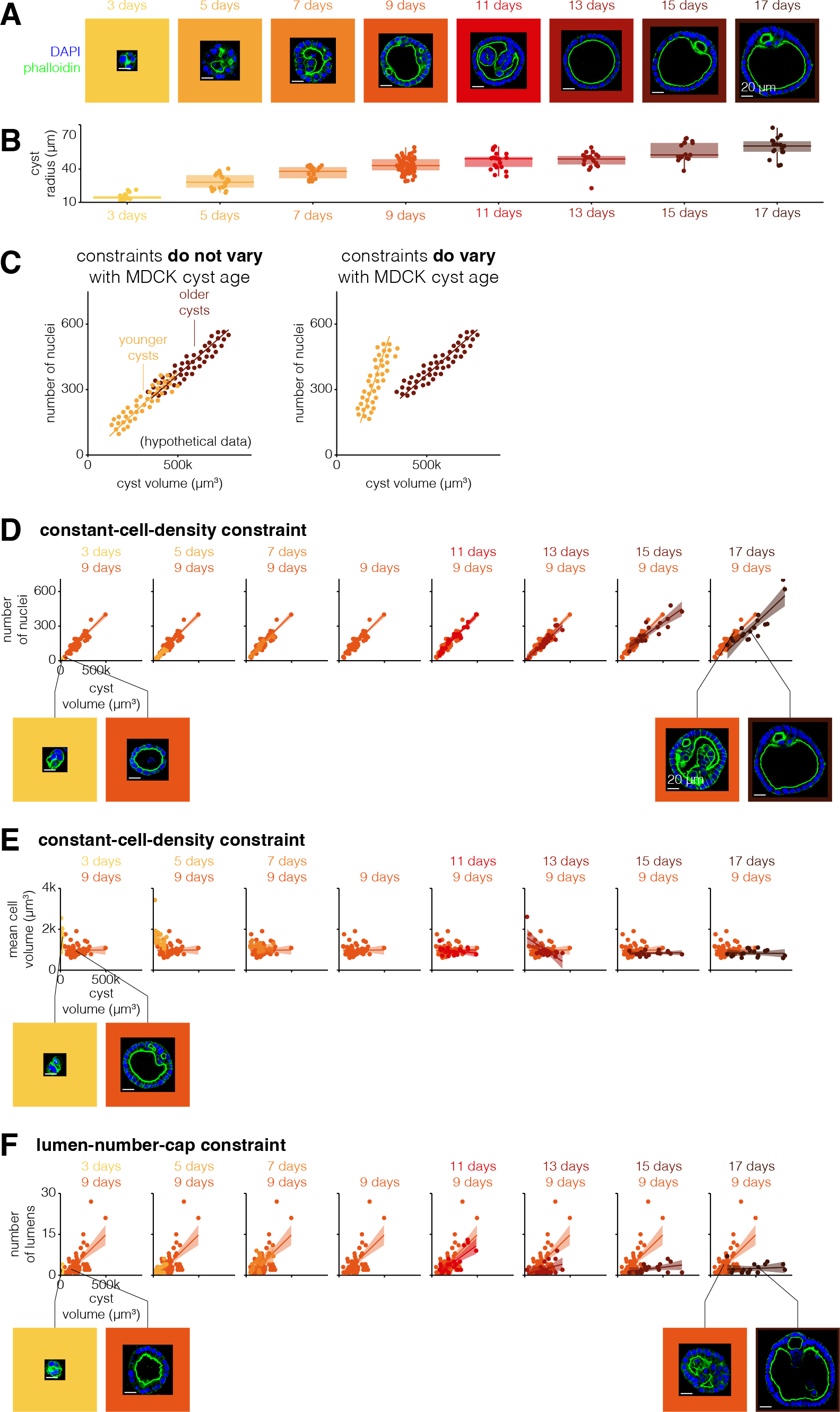
Constraints on MOCK cyst morphologies vary with age. **A.** Example MOCK cysts cultured for 3-17 days. **B.** Quantification of cyst radius for MOCK cysts of different ages. **C.** Example of a constraint that does or does not vary with MOCK cyst age for hypothetical data. **O.** Number of nuclei versus cyst volume for MOCK cysts of each age. Each age is represented by one color, and 9 day old MOCK cysts are repeated on each graph for reference. The line represents the line of best fit and the shaded area represents the 95% confidence interval. Example MOCK cysts with approximately the same number of nuclei but different volumes. **E.** Mean cell volume versus cyst volume for MOCK cysts of each age. Each age is represented by one color, and 9 day old MOCK cysts are repeated on each graph for reference. The line represents the line of best fit and the shaded area represents the 95% confidence interval. Example MOCK cysts with approximately the same volume but different mean cell volumes. **F.** Number of lumens versus cyst volume for MOCK cysts of each age. Each age is represented by one color, and 9 day old MOCK cysts are repeated on each graph for reference. The line represents the line of best fit and the shaded area represents the 95% confidence interval. Example MOCK cysts with approximately the same number of lumens but different cyst volumes.

We first wondered if the constant-cell-density constraint varied for cysts of different ages (Fig. 2D, Supp. Fig. 12). We found that cysts of all ages obeyed the constant-cell-density constraint on the number of nuclei and cyst volume. This relationship was confirmed by PCA run on the complete dataset including all time points, in which the principal components remained the same, but the principal component for cyst size accounted for more total variance due to size differences with age (Supp. Fig. 13). However, the exact nature of this constraint, specifically the slope and intercept of the linear relationship, varied with the age of the cyst. Cysts cultured for 3-5 or 13-17 days obeyed a constant-cell-density constraint with a smaller slope; i.e., they had fewer nuclei per cyst volume. As cysts age, they get larger, so in principle it could be that older cysts could have a lower cell density because all cells are larger, or it could be that there is a threshold volume beyond which cell density decreases. We found that amongst older cysts with smaller sizes (that matched those of middle-aged cysts), the densities were similar to those of middle-aged cysts, suggesting the latter threshold scenario (Fig. 2E, Supp. Fig. 14). Younger cysts, however, had a uniformly lower cell density than middle-aged cysts.

We wondered whether other constraints were similarly affected by age. We looked at the age dependence of the lumen-number-cap constraint (Fig. 2F, Supp. Fig. 15). We again found that cysts of all ages obeyed a constraint on the number of lumens per cyst volume. However, this constraint changed with age: we found that younger cysts cultured for 3 days had a higher maximum number of lumens per cyst volume, and cysts cultured for 13-17 days had a lower maximum number of lumens per cyst volume. The decrease in the number of lumens per cyst as cysts age beyond 9 days suggests that multiple lumens in a cyst are either merging or disappearing as cysts grow older. It is difficult to speculate what mechanisms might govern this age-dependent quantitative change in constraints without perturbations. However, given that very young cysts seem to obey an entirely different set of constraints than older cysts, it may be that some of these differences are due to the establishment of apico-basal polarity around the lumen considering the work of Vasquez et al (Vasquez et al. 2021). Also, we noticed in young cysts that the threshold of size and number of nuclei in order to form a lumen was 10,000 µm^3^ and 7 nuclei, respectively (Supp. Fig. 16). Together, our results point to constraints as being dynamic entities that can change as cysts grow and develop.

### Orug and environmental perturbations can change constraint parameters but do not break them

Having found constraints on MDCK cyst morphologies, we wondered whether these constraints applied to cysts whose morphology we perturbed using exogenous agents. For example, if we perturb the cysts with a drug which makes the cysts larger, will the same constant-cell-density constraint still apply? If not, there are two ways that the constraint could be disobeyed. One is that the perturbed cysts could follow the same constraint but with different parameters, for example, by changing the slope of the relationship between cell number and cyst volume. Alternatively, a perturbation could qualitatively remove a constraint and decouple, for example, the number of nuclei and cyst volume. Either of these possibilities would suggest that the constraints on MDCK cyst morphologies are context-specific.

There are few references to drugs which modify the morphology of MDCK cysts in the literature. Thus, to identify drugs that change MDCK cyst morphologies we designed a high-throughput drug screen of small molecule drugs from Selleck Chemicals Bioactive Compound library. This library contains -2,000 small molecule drugs including kinase and epigenetic inhibitors as well as ion channel, metabolic, and cancer compounds. To conduct the screen, we plated MDCK cells in 384-well plates, added 1 µM of each drug, and allowed the cells to grow into cysts for seven days, at which point the cysts were fixed and imaged. We then quantified the area of each cyst and the average across cysts for each perturbation. We found that, while most drugs did not appear to change the area of the cyst relative to controls, there were many drugs which made the cysts smaller or larger (Fig. 3A). We considered “hits for larger cysts to be the drugs that were on both the list of the top 100 drugs as ranked by fold change and the list of the top 100 drugs as ranked by z-score, a total of 78 drugs. We found 80 “hits” for smaller cysts using the same approach. We found that hits that resulted in smaller cysts were enriched for drugs in the kinase, epigenetic, and cancer categories, while hits for larger cysts were enriched for kinases, cancer, and G protein-coupled receptor categories (Supp. Fig. 17). To gauge how many of our hits may have arisen by chance, we screened a portion (1/7th) of the drug library again. We found that there was a correlation (r=0.63) between the fold change in cyst area from the screen and this targeted replication, suggesting that the majority of our hits were not random (Supp. Fig. 18). The screen hits represented potential candidates for perturbing MDCK cyst morphologies and then asking whether perturbed MDCK cysts obey the same constraints as unperturbed cysts.

**Figure 3:**
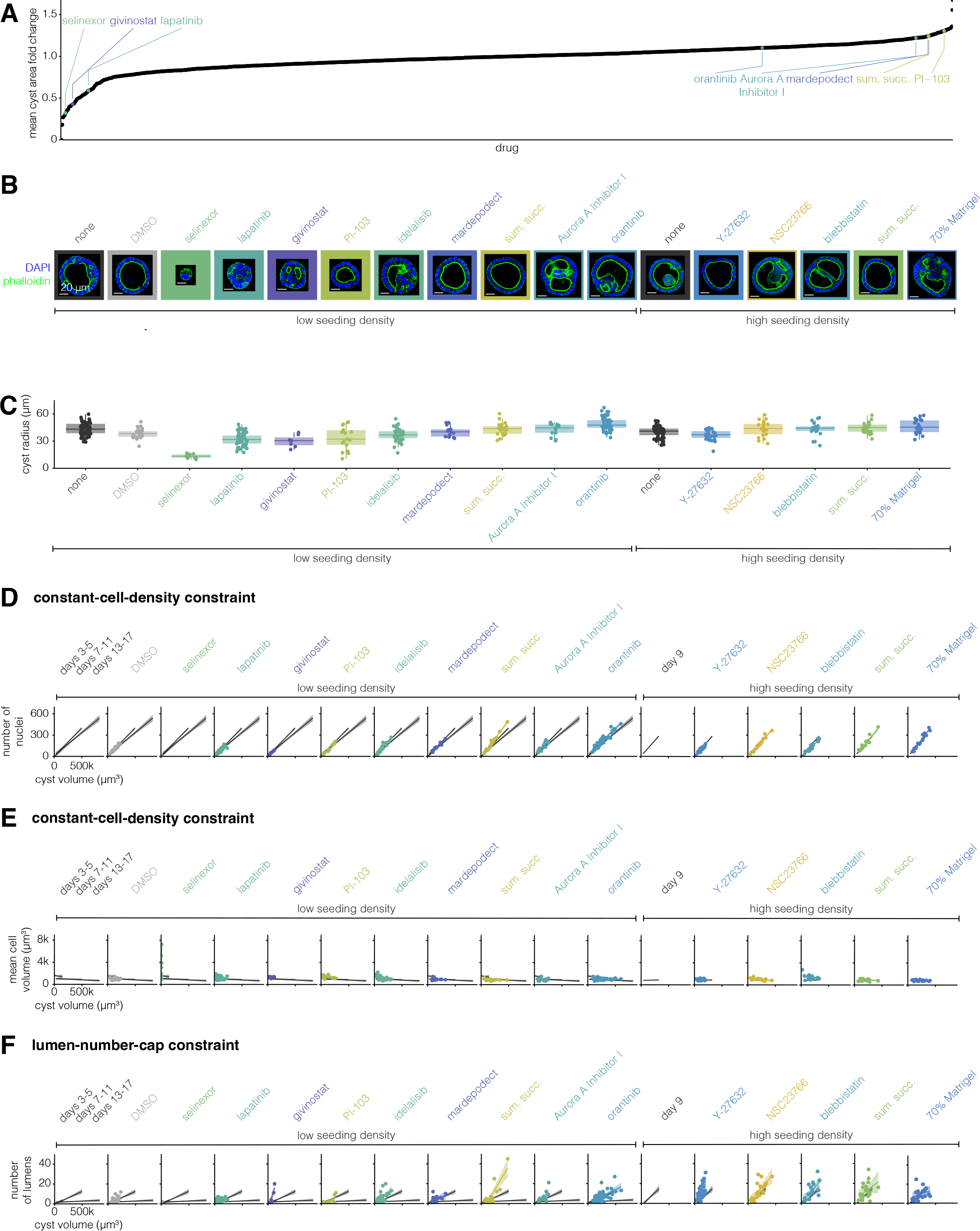
Drug and environmental perturbations can change constraint parameters but do not break them. **A.** Fold change in cyst area versus each drug from the screen. Annotated drugs are those used in further experiments. **B.** Example MOCK cysts for each perturbation. The seeding cell density, low (25,000 cells/mL) or high (100,000 cells/mL), and the drug added to the culture media are indicated below the image. The MOCK cyst shown is one with approximately average radius for that perturbation. **C.** Quantification of cyst radius for MOCK cysts exposed to different perturbations. **O.** Number of nuclei versus cyst volume for MOCK cysts exposed to different perturbations. Each perturbation is represented by one color. The line represents the line of best fit and the shaded area represents the 95% confidence interval. The line of best fit and 95% confidence interval for three groups of unperturbed MOCK cysts (3-5 days, 7-11 days, and 13-17 days) are shown in gray for reference. **E.** Mean cell volume versus cyst volume for MOCK cysts exposed to different perturbations. Each perturbation is represented by one color. The line represents the line of best fit and the shaded area represents the 95% confidence interval. The line of best fit and 95% confidence interval for three groups of unperturbed MOCK cysts (3-5 days, 7-11 days, and 13-17 days) are shown in gray for reference. **F.** Number of lumens versus cyst volume for MOCK cysts exposed to different perturbations. Each perturbation is represented by one color. The line represents the line of best fit and the shaded area represents the 95% confidence interval. The line of best fit and 95% confidence interval for three groups of unperturbed MOCK cysts (3-5 days, 7-11 days, and 13-17 days) are shown in gray for reference.

We further manually grouped hits for smaller and larger cysts according to their targets (Supp. Table 3-4). We selected four drugs from our list of hits from the screen that increased cyst size from groups targeting mammalian target of rapamycin, aurora kinase, phosphodiesterase, and serotonin. Similarly, we selected three drugs that made cysts smaller from groups targeting epidermal growth factor receptor, histone deacetylases, and exportin-1. Given the relatively small number of factors we were able to rigorously test, we did not perform enrichment analysis on the categories of factors that came up as targets of our drug screen. We additionally used the following perturbations we thought likely to change MDCK cyst morphology based on the literature: idelalisib, oratinib, Y-27632, NSC23766, and blebbistatin. We plated MDCK cells to form cysts, immediately added these drugs at a range of concentrations, and then grew the cysts for 9 days (Table 2). Additionally, we tested two non-drug perturbations, cell seeding density (by culturing MDCK cysts with a higher initial cell density) and dilute Matrigel. We then fixed, stained, and imaged the perturbed cysts as described above, after which we measured the same set of morphological features (Fig. 3B-C). We found that the screen hits that we expected to make cysts smaller did indeed lead to smaller cysts, but none of the ones predicted to make them larger did so. We found that increased seeding density nor dilute Matrigel had no effect on the size of the cysts.

Note that many of the hits from the screen targeted proliferation, suggesting that perhaps effects on size were a necessary consequence of changes to proliferative capacity. However, many other drugs in the screen also targeted proliferation but had no effect on cyst size, arguing against this possibility.

We then wondered whether perturbed cysts obeyed the same constant-cell-density and lumen-number-cap constraints as unperturbed cysts. We found that, with the exception of five drugs, perturbed cysts obeyed the same constant-cell-density constraint as unperturbed cysts (Fig. 3D-E). Cysts perturbed with sumatriptan succinate (a serotonin receptor inhibitor) or blebbistatin (a myosin II inhibitor) had more nuclei per given cyst volume than any age of unperturbed cysts. Cysts perturbed with selinexor (an exportin-1 inhibitor), Y-27632 (a ROCK inhibitor), NSC 23766 (a Rac inhibitor), or blebbistatin had larger nuclei per given cyst volume. Notably, the relationship between cell number, size, and cyst volume is still constrained for cysts perturbed with any of these drugs, but the parameters of the constraint (specifically the slope) are different from unperturbed cysts.

We then wondered how a perturbation which does change a constraint influences other constraints-if cysts perturbed with drug X do not obey the constant-cell-density constraint, must they also not obey the lumen-number-cap constraint? We found that cysts perturbed with givinostat (a histone deacetylase inhibitor), idelalisib (a phosphoinositide 3-kinase delta isoform inhibitor), sumatriptan succinate, Aurora A Inhibitor I, Y-27632, and blebbistatin had more lumens in a given cyst volume than unperturbed cysts of any age (Fig. 3F). Thus, cysts perturbed with these drugs do not obey the same lumen-number-cap constraint of unperturbed cysts, instead they obey a constraint with a larger slope. We found that some perturbations (selinexor, givinostat, idelalisib, Aurora A Inhibitor I, and NSC 23766) changed only one constraint, but others (sumatriptan succinate, Y-27632, and blebbistatin) changed both. Given the targets of these perturbations, we can hypothesize as to which mechanisms may regulate these constraints. We saw that exportin 1 was important for the constant-cell-density constraint, HDAC, PI3K, and Aurora A kinase for the lumen-number-cap constraint, and serotonin receptors were important to both constraints. We similarly found some mechanisms perturbed MDCK cysts along a single axis in principal component space, while others perturb cysts along two axes (Supp. Fig. 19), suggesting that the axes of MDCK cyst variation may not be governed by independent mechanisms. These factors all work through quite different pathways; however, given the relatively small number of perturbations we analyzed, it was difficult to uncover any general rules on which pathways affected which constraints. In combination, this suggests that the set of morphologies available to MDCK cysts is richer than unperturbed cysts would suggest.

### Constraints of perturbed cysts do not add together or average out when multiple perturbations are applied

Exposing MDCK cysts to hepatocyte growth factor (HGF) in collagen gels induces cells to send out spindly extensions (Montesano, Schaller, and Orci 1991; Montesano et al. 1991). These extensions form the groundwork for a chain of cells to proliferate and ultimately form tubules. We wondered how such a perturbation might change the constraints on cyst morphologies we identified, given that it leads to known, qualitative changes in cyst morphology. We added HGF to MDCK cysts as they grew for nine days (Fig. 4A). We used our imaging and feature quantification approach to characterize the constraints on cysts exposed to HGF. We asked whether our measurements of cyst shape captured the morphology of extensions in HGF-perturbed cysts. We found that one of the primary differences between unperturbed and HGF-perturbed cysts was cyst solidity, a measure of convexity (cysts with more involutions or protrusions have lower solidity than circular or elliptical cysts) (Fig. 4B, Supp. Fig. 20). Mean cyst solidity decreased from 0.93 for unperturbed cysts to 0.75 for HGF-perturbed cysts. We also found that cysts exposed to HGF were larger, on average, than unperturbed cysts (Fig. 4C).

**Figure 4:**
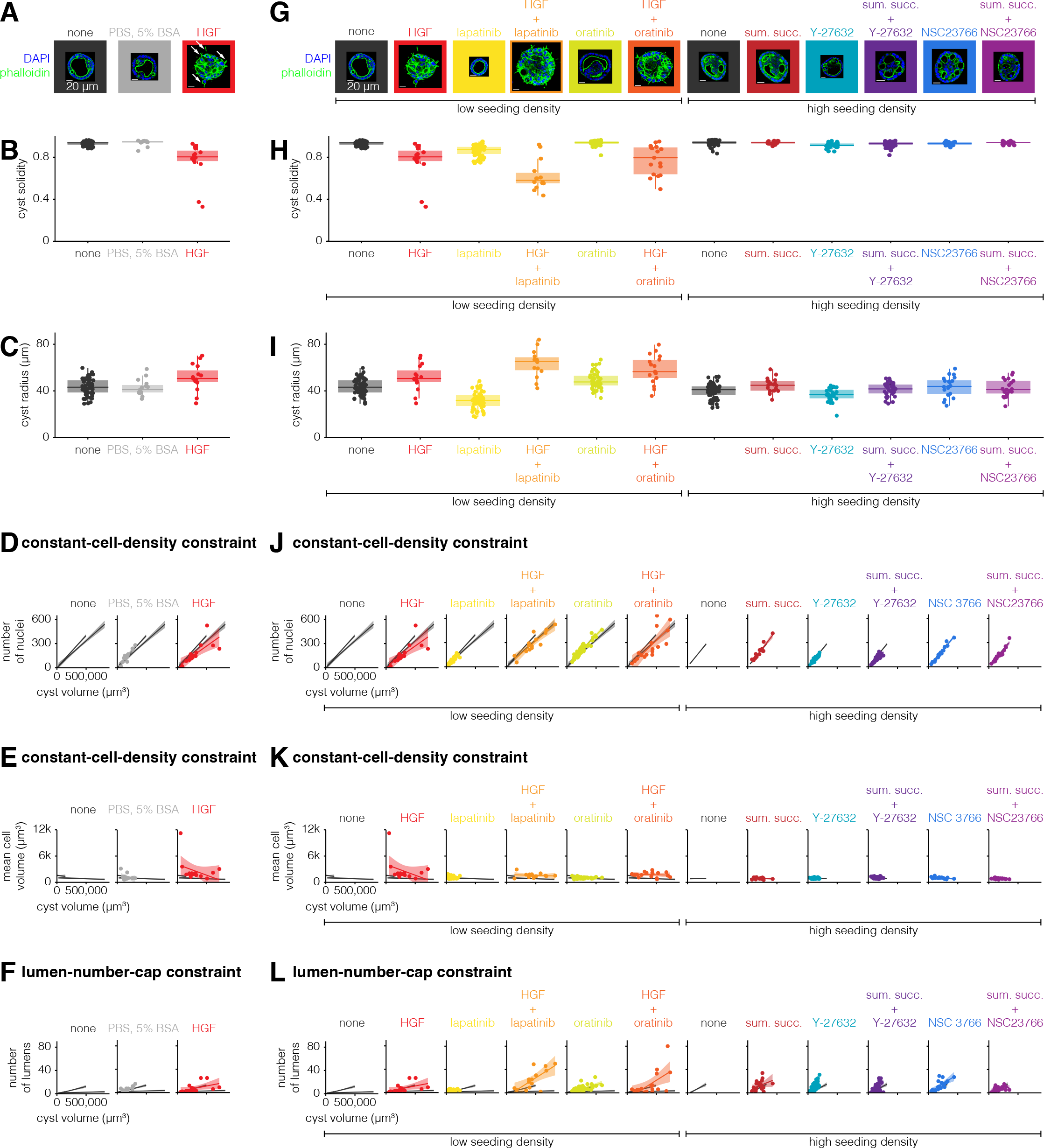
Constraints of perturbed cysts do not add together or average out when multiple perturbations are applied. **A, G.** Example MOCK cysts cultured for each perturbation. The seeding cell density, low (25,000 cells/ml) or high (100,000 cells/ml), and the drug added to the culture media are indicated below the image. The MOCK cyst shown is one with approximately average radius for that perturbation. White arrows indicate spindle-like extensions. **B, H.** Quantification of cyst solidity for MOCK cysts exposed to different perturbations. **C, I.** Quantification of cyst volume for MOCK cysts exposed to different perturbations. **D, J.** Number of nuclei versus cyst volume for MOCK cysts exposed to different perturbations. Each perturbation is represented by one color. The line represents the line of best fit and the shaded area represents the 95% confidence interval. The line of best fit and 95% confidence interval for three groups of unperturbed MOCK cysts (3-5 days, 7-11 days, and 13-17 days) are shown in gray for reference. **E, K.** Mean cell volume versus cyst volume for MOCK cysts exposed to different perturbations. Each perturbation is represented by one color. The line represents the line of best fit and the shaded area represents the 95% confidence interval. The line of best fit and 95% confidence interval for three groups of unperturbed MOCK cysts (3-5 days, 7-11 days, and 13-17 days) are shown in gray for reference. **F, L.** Number of lumens versus cyst volume for MOCK cysts exposed to different perturbations. Each perturbation is represented by one color. The line represents the line of best fit and the shaded area represents the 95% confidence interval. The line of best fit and 95% confidence interval for three groups of unperturbed MOCK cysts (3-5 days, 7-11 days, and 13-17 days) are shown in gray for reference.

Given the qualitative difference in morphology of HGF-perturbed cysts, we wondered if HGF-perturbed cysts obeyed the constraints obeyed by unperturbed MDCK cysts. We found that HGF-perturbed cysts did not obey both aspects of the constant-cell-density constraint: while cysts perturbed with HGF had the same number of cells per cyst volume, the cells were larger than those of unperturbed cysts of any age (Fig. 4D-E). How do HGF-perturbed cysts have the same number of cells per cyst volume, but larger cells than unperturbed cysts? One possibility was that HGF-perturbed cysts have a smaller proportion of their volume taken up by lumens and a larger proportion of the volume occupied by cells. Interestingly, HGF-perturbed cysts do obey the lumen-number-cap constraint, suggesting that what lumens HGF-perturbed cysts have are smaller in size but similar in number (Fig. 4F). The smaller proportion of volume taken up by the lumens could result from cells being taller or adopting a different configuration. We found that the cells often formed multi-cell layers, which allows for larger cells to occupy the same organoid volume while maintaining the same total number of cells per volume. It also suggests that the strict proportionality between cell number and organoid volume is maintained despite disruptions to cellular configurations and hence cell number may not be controlled by morphology *per se*.

Given that HGF qualitatively changed some features of MDCK cysts, we wondered what the morphological effects would be upon combining HGF with the previously used perturbations that engendered more quantitative changes. For example, would a perturbation that produces cysts with extensions (HGF) and a perturbation that produces smaller cysts yield small cysts with extensions? We perturbed MDCK cysts for nine days with either HGF alone or HGF in combination with lapatinib or orantinib (Fig. 4G). To ensure that any differences we noticed could not be explained by possible cross-talk between HGF and EGFR (Jo et al. 2000), we additionally perturbed MDCK cysts with sumatriptan succinate alone or in combination with Y-27632 or NSC23766. We found that cysts exposed to HGF and lapatinib or oratinib had lower solidity than cysts exposed to only one of these perturbations (Fig. 4H). We found that cysts exposed to HGF, alone or in combination, were also larger, on average, than control cysts (Fig. 4I). Taken together, the morphological changes induced by HGF and another perturbation suggest that the effects of individual perturbations do not necessarily combine additively when administered simultaneously.

We then wondered how the constraints of cysts perturbed with one drug changed when the cysts were exposed to a second drug. One possibility is that doubly-perturbed cysts obeyed a set of constraints that averaged the constraints obeyed by singly-perturbed cysts (assumption of linearity). Another possibility is that doubly-perturbed cysts obeyed the same set of constraints as only one of the perturbations, suggesting that some drugs may be able to override the effects of others, or some other non-linear interaction. One might expect that linearity would hold for small doses of drug, but that nonlinear aspects of the regulatory processes may appear for larger doses. We found that sometimes one perturbation overrode the effects of the other and that sometimes doubly-perturbed cysts did not obey the same constraints that the singly-perturbed cysts did (Fig. 4J-L). Perturbations overrode the effects of one another in many combinations and for both constraints. We observed that while HGF alone obeyed a different constant-cell-density constraint, when used in combination with either lapatinib or oratinib the cysts obeyed the same constraint as unperturbed cysts. We likewise found that the effects of Y-27632 on the lumen-number-cap constraint and the effects of NSC23766 on the constant cell density constraint were both canceled out by sumatriptan succinate. We additionally found many examples where singly-perturbed MDCK cysts obeyed the same constraints, but when those perturbations were used in combination the cysts did not obey the same constraints. We found that while neither HGF, lapatinib, nor oratinib alone changed the lumen-number-cap constraint, cysts perturbed with both HGF and lapatinib or oratinib had higher lumens per cyst volume (Fig. 4J-L). We also found this to be true for sumatriptan succinate and Y-27632 for the constant-cell-density constraint. In totality, the many differences between the constraints obeyed by double-perturbed cysts and single-perturbed cysts suggests that the effects of any given perturbation do not appear to simply add together, but rather can combine in unanticipated ways.

It is important to note that many of the perturbations affected particular classes of biological processes, such as proliferation (e.g. lapatinib, oratinib, etc) and those affecting morphological processes (e.g. HGF, Y-27632, etc). It is possible that the use of different combinations with drugs perturbing the same processes could have more predictable effects.

### Enteroids show a similar set of perturbable constraints

The MDCK cyst formation system is relatively simple (spherical shape, single cell type), and so we wondered whether our approach would apply to more complex organoid systems. We chose to apply our approach to the enteroid system, which is an organoid system mimicking intestinal crypt and villus structures. In this system, enteroids can be derived from a single intestinal epithelial stem cell, from which multiple cell types emerge to form a structure that in some ways reflects the crypt and villus structure of the intestinal epithelium. In that structure, stem cells are at the base of the crypt and give rise to the other cell types, including Paneth cells, goblet cells, enterocytes, transit amplifying cells, and enteroendocrine cells (Fig. 5A). In (Lukonin et al. 2020), the authors performed high-throughput quantitative measurements on enteroids exposed to a wide variety of perturbative conditions (2,455 conditions), generating a dataset that matched many of the criteria for our approach. Their measurements included enteroid area and a number of associated morphological quantities (Fig. 5B). Furthermore, they performed antibody staining for markers of Paneth cells and enterocytes, along with nuclear staining with DAPI, yielding the fluorescence intensity of these markers as a proxy for the number of cells of these types (and total cells in the case of DAPI). It is worth noting here that while the intensities serve as a proxy for cell number, they do not give the relative location of these cell types, which are data that could be useful in future studies.

**Figure 5:**
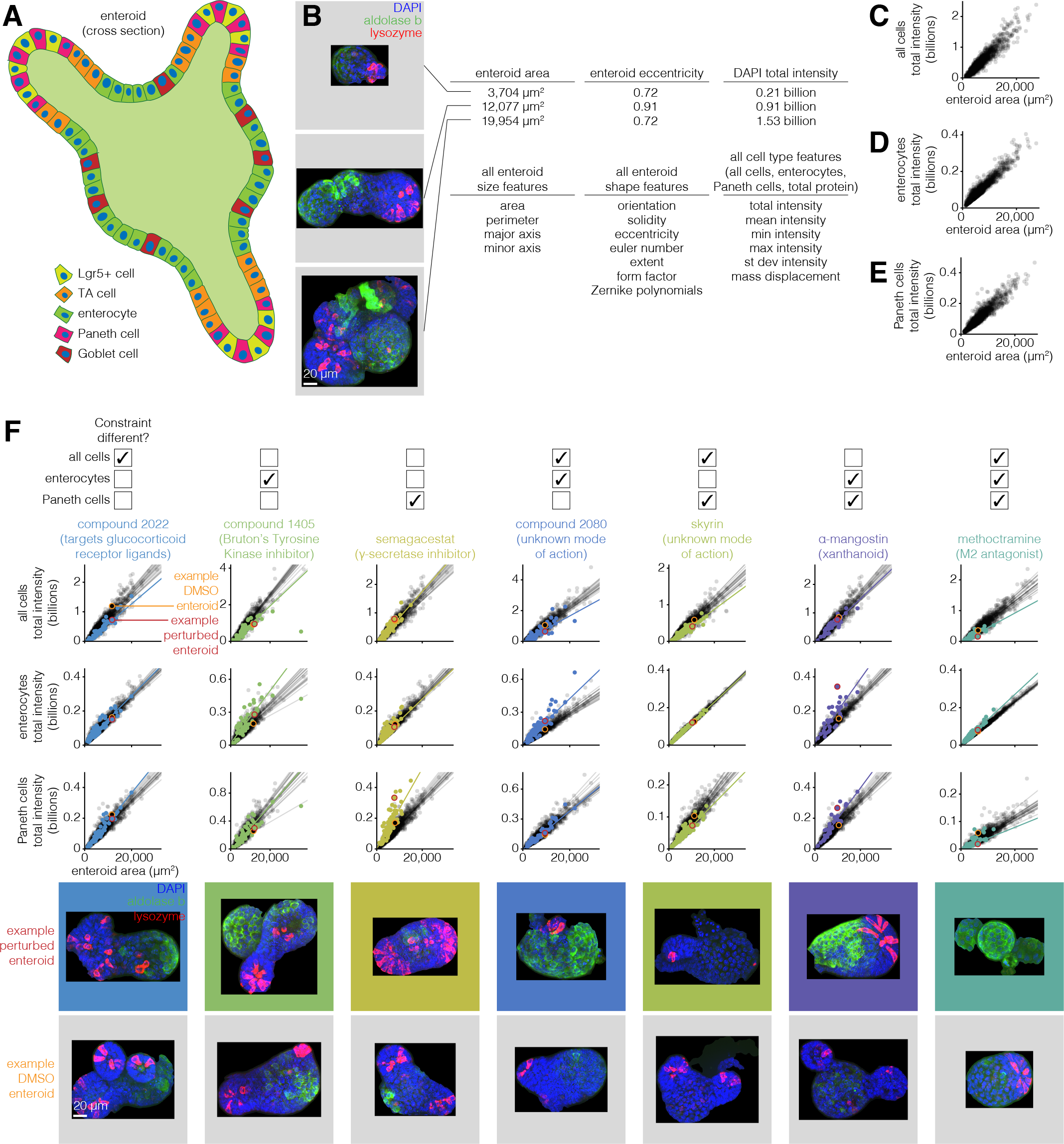
Enteroids obey a similar set of constraints that can also be modulated by drugs. **A.** Schematic of enteroid with different cell types identified. **B.** Example enteroids with different morphological and cell type features. **C.** Total intensity of all cells versus cyst volume for unperturbed enteroids. **D.** Total intensity of enterocytes versus cyst volume for unperturbed enteroids. **E.** Total intensity of Paneth cells versus cyst volume for unperturbed enteroids. **F.** Constraints from C-E for enteroids perturbed with drugs. An example perturbation is shown for each set of constraints that can be different. Unperturbed enteroids are plotted in gray and perturbed enteroids are plotted in color. The lines represent the line of best fit and are plotted for each well that the enteroids were cultured in. The enteroids in the images are circled on the graph (unperturbed enteroid in orange and perturbed enteroid in red).

We first performed PCA on the features from this enteroid dataset to look for constrained covariation between features (Supp. Fig. 21). We found that there were several principal components that appeared to explain a significant fraction of the variance. The first principal component appeared to correspond to size, and again revealed a strong correlation between size and total number of cells as well as the number of enterocytes and Paneth cells. Looking at the second and third principal components, the density (or mean intensity) of Paneth cells appeared to correlate with protein density and density of cells independently of any enterocyte features (PC2). However, the density of enterocytes seemed to correlate with protein density independent of all cell and Paneth cell features (PC3). Together, the PCA revealed a number of axes of variation with respect to the relative numbers of different cell types.

We first wanted to establish that the scaling properties that we observed in MDCK had some analogy in the enteroid system (Fig. 5C-E). We found that all three fluorescence intensity quantities (DAPI/number of nuclei, aldolase b/number of enterocytes, lysozyme/number of paneth cells) scaled strongly with enteroid area, showing that the numbers of each of these cell types in general scaled with the total size of the enteroid. (Note that the nature of these data was rather different than that collected for MDCK, so we performed a different set of quality control steps; see methods for details.)

With these strong linear relationships in hand, we went about searching through the set of perturbations looking for examples of perturbations that altered these relationships and combinations thereof. We looked for perturbations that gave large fold changes to the slope or intercept of a linear fit as compared to unperturbed enteroids, with the further criterion that the majority of replicates of the drug (both replicates when there were two total replicates, or at least two out of three if there were three total replicates) must show the change (Fig. 5F). We found 118 perturbations that caused a change in at least one of the constraints. We confirmed these changes by looking at the raw images, which demonstrated that the changes in fluorescence intensity resulted from changes in the number of cells of the various types predicted (as opposed to changes in staining intensity of the same number of cells). Indeed, for the seven possible combinations of constraint changes (changes in 1, 2 or all 3 of the constraints at once), we found at least one perturbation (often several) for each constraint change combination. Our results show that the scaling property of the number of cells of a particular type with organoid size are maintained across perturbations, but that the scaling constant itself (i.e., the proportion of cells of a particular type out of the total) are subject to change.

In conclusion, we found that our approach was able to identify constraints in a more complex organoid system, and that these constraints were perturbable. It is important to note that the set of features in these data is likely far richer than what was currently captured in our measurements; for instance, quantities associated with the distances between various cell types were not computed. A richer set of features may lead to a richer set of constraints.

## Discussion

Here, we sought to quantify constraints on MDCK cyst morphologies. We found the MDCK cysts obey a number of constraints, and that the majority of their morphological variation can be explained by three dimensions. We also found that some constraints on MDCK cyst morphologies vary with age and perturbations.

It remains unclear what underpins the constraints on cyst (or, more generally, organoid) morphologies. One could imagine any number of potential mechanisms, any one of which might be critical to a constraint by itself or in combination with many others. Such mechanisms may be based upon conventional biochemical signaling (such as signaling between cells to control proliferation), or may involve mechanical sensing of variables such as membrane curvature. A related question is whether these mechanisms map in a one-to-one manner onto each of the principal components of variation we detected. I.e., is there one mechanism that governs cell size and related variables, and another mechanism that governs lumen number? In principle, there is no need for any one mechanism to map in such a direct way onto a particular axis of variation. Our data with various drugs suggests that it is entirely possible for a mechanism to affect just one or multiple principal components, so there is no reason to predict that individual mechanisms would be restricted to affecting just a single axis.

While many potential mechanisms may be compatible with our experimental data, perturbations will be required to exclude certain classes of models and establish causality. Such molecular mechanisms, if identified, are enormously powerful and are a critical ultimate goal for molecular biology. Given the relatively small number of perturbations, it was difficult to connect pathways with phenotypes. It is possible that more extensive sets of perturbations may ultimately reveal the underlying molecular mechanisms responsible for particular constraints. However, it is also possible that the complexity of the underlying molecular pathways is too great and multi-faceted to ever fully relate to these constraints in an easily understood manner (Mellis and Raj 2015). Nevertheless, these constraints and others like them may constitute an effective “grammar” of organoid morphology that one may be able to build upon irrespective of the molecular details. Knowledge of the building blocks of organoid morphologies may also inform generative quantitative models of morphogenesis that could be evaluated for their ability to recapitulate such constraints.

We also found that while some perturbations altered cyst parameters within constraints, others changed the nature of the constraint. Knowledge of which types of perturbations lead to which type of effect might aid in the development of an instruction manual for building designer organoids, potentially existing in very different parts of parameter space than normal organoids. Our framework may reveal the parameters one may be able to manipulate organoids using the rules learned by these systematic perturbations. Such organoids may have properties that make them more useful for particular applications in regenerative medicine or as disease models. It may also be possible, in principle if not in practice, to destroy a constraint with sufficient perturbation. For example, the right perturbation might completely decouple cyst volume from the number of cells. Future work could search for perturbations with such effects by combining high throughput drug screens with our detailed quantification of organoid morphologies. With the ability to decouple morphological constraints, we might be able to engineer organoids to adopt entirely novel configurations.

One principal technical challenge in the scaling of approaches such as the one we took here is the extraction of annotations of MDCK cyst structures from microscopy images. Our assumption was that we would need highly accurate annotations to reveal subtle constraints on MDCK cyst morphologies, and those annotations proved difficult to fully automate. For this reason, we chose to build an interface that enabled us to manually correct annotations from any algorithm. Our hope is that this approach and the software is of use to others looking to annotate images, structures, or tissues for which automated solutions have yet to be developed. Deep learning has produced great advances in automatic image segmentation (Moen et al. 2019), and it is possible that the application of these methods, once fully automated and of very high quality, would allow us to obtain much larger numbers of annotations compared to our combination of automated algorithms and manual annotation review. It is also worth considering what level of segmentation accuracy is needed for the question at hand. Future work that quantifies what degree of segmentation accuracy is needed for a given question may guide efforts to develop segmentation algorithms. In addition, it is possible that alternative strategies for culturing or imaging the cysts might make the images easier to segment. One such example would be to plate all MDCK cysts at the same distance from the bottom of the well. Future work could also use the large and highly accurate segmentations produced in this work to train deep learning models specific to this task.

We focused primarily on MDCK cysts for our proof of concept because of their simplicity, both morphologically and in terms of the number of cell types involved (in this case, just one cell type). We also applied our methodology to the more complex enteroid system that has several cell types that interact in various ways; however, the feature set available for each enteroid was relatively less rich due to the complexities of quantifying those features in a complex three dimensional set of cells. We have demonstrated, in principle, that our approach can determine the ways in which organoid morphologies are either constrained or free to vary.

However, it is important to note that more complex organoid systems may have higher degrees of dimensionality and non-linearities that may require more sophisticated approaches and analyses. It will be interesting in the future to apply this framework to such organoids to see what constraints are obeyed, as well as to map the relationship between an organoid’s morphological constraints to an organoid’s functional characteristics.

## Materials and Methods

### MOCK Cyst Culture

Madin-Darby Canine Kidney-II cells (MilliporeSigma, 00062107-1VL) were maintained by culturing them in 2D on traditional 10 cm cell culture-coated dishes (Corning, 353003). The media for both the adherent 2D cells and cysts was MEM media (MediaTech, MT10-010-CM) with 10% Fetal Bovine Serum (Fisher, 16000044) and 1X Penicillin-Streptomycin (Invitrogen, 15140122). When the cells were between 30-70% confluence there were dissociated to make cysts. The cells were briefly washed with 5 mL of DPBS (Gibco, 14190136). Then, 1 mL of 0.25% Trypsin-EDTA (Gibco, 25200056) was added and the plate was incubated at 37 °C, 5% CO_2_ for 10-15 minutes. The trypsin was inactivated with 9 mL of media and the solution was pipetted over the dish three times to ensure all cells detached. The cells were pelleted for 2 minutes at 1000 rpm and then suspended in 500 µL to 1 mL media. The cell concentration was quantified using a BioRad TC20 Automated Cell Counter. The cells were added to ice-cold thawed Matrigel (Corning, 354234) at a concentration of 25,000 cells/mL. The middle of a well of a Nunc Lab-Tek 8-well Chambered Coverglass (Fisher, 12-565-470) was coated with 5 µL of pure Matrigel. Then, 25 µL of the cell-Matrigel suspension was overlaid on top of the coating. The chamber was incubated at 37 °C, 5% CO_2_ for 30 minutes to solidify the Matrigel. Then, 200 µL of media was added on top of the solidified Matrigel. The cysts were returned to the 37 °C, 5% CO_2_ incubator and cultured for 3-17 days. The media was replaced every other day.

### Imaging

MDCK cyst fixation and staining was performed at room temperature with two brief washes with 1X PBS (Ambion, AM9624) between each step. When the cysts were ready to be imaged they were fixed in their culture chambers with 1.85% formaldehyde (MilliporeSigma, F1635-500ML) in PBS for 30 minutes. They were permeabilized with 0.1% Triton X-100 (MilliporeSigma, T8787-100mL) in PBS overnight. The cysts were then blocked with 5% Bovine Serum Albumin (MilliporeSigma, A7906-100G) in PBS for 1 hour. The cysts were then incubated with 1:15 488-phalloidin (Invitrogen, A12379) and 1:30 DAPI (Fisher, D3571) in PBS for at least 6 hours before imaging. The cysts were imaged on a Zeiss Laser Scanning 710 Confocal Microscope using a 40X objective (Zeiss, water immersion, 1.1 NA, long working distance, LD C-Apochromat), 405 nm diode laser (Zeiss), and 488 nm argon-ion laser (LASOS). Each cyst was imaged from the bottom to a depth clearly beyond the middle point of the cyst. Cysts that were too far from the glass to image that deeply were not imaged.

### Morphological Quantification from Images

We wrote a custom MATLAB pipeline (https://github.com/arjunrajlaboratory/organoids2) to measure cyst morphological features from microscopy images by 3D segmenting the boundaries of the whole cyst and each of its lumens and nuclei. To segment the cyst and lumen boundaries our general approach was to guess the boundary on each image slice using the phalloidin image and then manually correct the boundary as needed. To guess the cyst boundary on each slice we set an empty corner of the image as the starting boundary and expanded that boundary outward until the intensity of those pixels was above a user-defined threshold. We applied the same approach to guess the lumen boundaries, except we identified the starting point as the largest object after the slice had been processed with a Laplacian of Gaussian edge detector. We then manually reviewed these 2D boundaries and corrected them as needed (Supp. Fig. 2). Once these 2D boundaries were finalized they were combined to form 3D boundaries. We obtained 3D cyst boundaries by assuming all 2D cyst annotations belonged to the same object. We obtained 3D lumen boundaries by computationally identifying which boundaries touched one another when stacked in 3D. To 2D segment the nuclear boundaries we used cellpose to segment the nuclei on the original image slices. We also sliced the image stack orthogonally from its original slicing, such that moving from slice-to-slice moves left-right across the cyst, rather than up-down. We also used cellpose to segment the nuclei on these orthogonal slices. We then used the orthogonal 2D segmentations to guess which original 2D segmentations were connected to one another. We then manually reviewed these 3D connections and corrected them as needed (Supp. Fig. C).

Once we had 3D boundaries for the cyst, lumens, and nuclei, we wrote custom analyses to measure morphological features of size, shape, and number for each cyst (Supp. Table 1). For cysts with multiple lumens, we took the mean across all lumens. For cysts with multiple nuclei, we took both the mean and the standard deviation across all nuclei.

### PCA and Linear Models

In order to run PCA we first standardized the units of our features. We took the cube root of all volume features, the square root of all surface area features, and the inverse of the number of lumens. We then z-score normalized each feature. We ran PCA using the prcomp function from the R’s stats package (https://www.rdocumentation.org/packages/stats/versions/3.6.2). To estimate how much variance we could expect to be explained due to chance, we also ran PCA on randomized data. To randomize the data, we shuffled each column of a table where each row represents one cyst/enteroid and each column represents one morphological feature.

We fit linear models to various pairs of morphological features using ggplot2 (https://ggplot2.tidyverse.org/) and R’s stats package.

### Orug Screen

We first established MDCK-II cells with stable integration of GFP nuclear and mCherry cell membrane markers. The day before we planned to transfect the cells we plated them so that the cells would be -80% confluent at the time of transfection. The cells were cultured in media without antibiotics. The following day, we used Lipofectamine 2000 (Invitrogen, 11668019) to transfect the cells with H2B-GFP plasmid (https://www.addgene.org/11680/). Two days after transfection, we replaced the media with media containing penicillin, streptoymycin, and G418 (Mediatech, MT30-234-CR). We changed the media every other day. One week after transfection we single cell bottlenecked the cells. We then followed the same approach to transfect the cells with mem-mCherry plasmid (https://www.addgene.org/55779/).

To conduct the drug screen, we used Matrix WellMate to plate Matrigel with 35,000 cells/mL into 384-well plates. We then centrifuged the plates at 300 rpm for 1 minute to ensure the Matrigel-cell suspension fell to the bottom of the well. We polymerized the Matrigel by placing the plates in a 37 °C, 5% CO_2_ incubator for 30 minutes. We then added 20 µL of media with 20 mM HEPES and drug using a Perkin Elmer Janus Modular Dispensing Tool. The cysts were cultured for 7 days at 37 °C, 5% CO_2_.

To image the cysts, we fixed them with 20 µL of 8% formaldehyde for 30 minutes at room temperature. We washed the plates with PBS and then stained them with 1:2500 Hoescht in PBS overnight. We used a Molecular Device’s ImageXpress Micro XLS Widefield High-Content Analysis System to image each plate at 10X. We took 4 images, each at the height determined by the autofocus software, per well.

We then quantified the effect of each drug on cyst size using custom MATLAB scripts. First, we combined the three image channels. We then Gaussian filtered the image and binarized it using Otsu’s method. We then obtained the boundary of cysts by obtaining the boundary of all objects in the binary image that were bigger than 50 pixels and smaller than 1500 pixels. We calculated the area of each cyst using MATLAB’s regionprops function. We then calculated the average fold change in cyst area for each drug by dividing the average cyst area for the drug by the average cyst area for all control cyst from the same plate. We similarly calculate the z-score for each drug.

To identify hits that made the cysts larger, we found drugs in common between the list of top 100 drugs by fold change and the list of top 100 drugs by cyst area. To identify hits that made the cyst smaller, we found drugs in common between the list of both 100 drugs by fold chance and the list of bottom 100 drugs by cyst area.

### MOCK Cyst Perturbation Experiments

MDCK cysts were cultured using the above technique with the following exceptions. For drug perturbations, cysts were cultured in media containing drug throughout their entire growth (Supp. Table 2). Media was replaced every other day. For the high cell density perturbation, the cysts were plated from a cell-Matrigel suspension containing 100,000 cells/mL. For the 70% Matrigel perturbation, a solution of 70% Matrigel diluted with MDCK media was used to both coat the plate and culture the cysts. Cysts were fixed and imaged on the 9th day using the protocols described above. Note that in the drug screen the cysts were fixed on the 7th day (because the screening facility was concerned that longer timepoints would risk media evaporation). However, to more easily compare the effects of drug perturbations with age perturbations we decided to use the same reference age (9 days).

### Enteroid Analysis

Data on unperturbed and perturbed enteroids was provided by Ilya Lukonin and Prisca Liberali. Details on enteroid generation, culture, imaging, and image analysis can be found in their 2020 publication in Nature.

### Oata and Code Availability

Our MATLAB pipeline for quantifying morphological features from microscopy images can be found at https://github.com/arjunrajlaboratory/organoids2. We additionally used MATLAB code from https://github.com/arjunrajlaboratory/rajlabimagetools.

All data and remaining code used for these analyses can be found at https://www.dropbox.com/sh/f2hq7n49nwkjjba/AABDe7hVJvisaGKF7t4hsZEha?dl=0. We used MATLAB to format images for all figures. All other analyses were done in R. We used a selection of color-blind friendly colors for figures 2 - 4 from https://personal.sron.nl/-pault/.

## Acknowledgements

We thank members of the Raj and Lengner labs for many useful discussions and suggestions. We thank Phil Burnham, Ian Mellis, and Karun Kiani for critical review of the manuscript. We also thank Benjamin Lamarck Emert, Phil Burnham, Yogesh Goyal, Amy Azaria, Karun Kiani, and Ian Dardani for help with fixing 3D annotations. We also thank Reka Hollandi from the Horvath lab for assistance in setting up and running their segmentation software, NucleAIzer. We also thank the Penn High Throughput Screening Core (Sara Cherry, David Schultz, and Christopher Rainville) and Penn Cell and Developmental Biology Microscopy Core (Andrea Stout, Jasmine Zhao) for their help and input. LEB acknowledged support from NIH T32 EB009384. AJH acknowledges support from NIH R35 GM133380, NIH R21 GM132831, and the Penn Institute for Translational Medicine and Therapeutics Pilot Grant. ZJG acknowledges support from the Chan Zuckerberg Biohub Investigator Program, NIH U01 CA199315, and the University of California, San Francisco Center for Cellular Construction DBI-1548297. AR acknowledges support from NIH R01 CA238237, NIH Director’s Transformative Research Award R01 GM137425, NIH R01 CA232256, NSF CAREER 1350601, NIH P30 CA016520, NIH SPORE P50 CA174523, NIH U01 CA227550, NIH 4DN U01 HL129998, NIH Center for Photogenomics RM1 HG007743, Chan Zuckerberg Initiative Human Cell Atlas 182724, and the Tara Miller Foundation.

## Author Contributions

LEB conceived of the project, performed the experiments, analyzed the data, and wrote the paper. JL, CC, and MCD all provided assistance with manual image annotation. NS and MKC performed some preliminary data analysis. IL and PL provided enteroid data and helpful input on the analysis of such data. AJH, JDM, and ZJG assisted with crowdsourced image segmentation and also provided valuable intellectual input on the conception of the project. AR conceived of the project, provided guidance in data analysis, and wrote the paper.

---

**Supplemental Table 1:** MDCK Cyst Morphological Features.

| Feature | Quantities Measured | Method |
| --- | --- | --- |
| volume ($\mu\text{m}^3$) | cyst volume<br>mean lumen volume<br>mean nuclear volume<br>standard deviation nuclear volume | 1. Calculate the number of voxels inside the object.<br>2. Multiply by the voxel volume. |
| cell volume ($\mu\text{m}^3$) | mean cell volume | 1. Subtract the volume of all lumens from the cyst volume.<br>2. Divide by the number of nuclei. |
| fraction of cyst volume | lumen fraction of cyst volume<br>nuclear fraction of cyst volume | 1. Sum the volume of all lumens/nuclei.<br>2. Divide by the cyst volume. |
| surface area ($\mu\text{m}^2$) | cyst surface area<br>mean lumen surface area<br>mean nuclear surface area<br>standard deviation nuclear surface area | 1. Use MATLAB's regionprops function to calculate the perimeter of the object on each image slice.<br>2. Sum the perimeters over all image slices.<br>3. Multiply by the size of the voxel in XY and the size of the voxel in Z. |
| XY radius ($\mu\text{m}$) | cyst XY radius<br>mean lumen XY radius<br>mean nuclear XY radius<br>standard deviation nuclear XY radius | 1. Calculate the image slice where the object has the largest area.<br>2. Calculate the center of the object on this slice.<br>3. Measure the distance between all boundary points (on this slice) and the center.<br>4. Take the mean. |
| Z radius ($\mu\text{m}$) | cyst Z radius<br>mean lumen Z radius<br>mean nuclear Z radius<br>standard deviation nuclear Z radius | 1. Calculate the maximum z coordinate of the object.<br>2. Subtract the minimum z coordinate of the object. |
| Z radius:XY radius | cyst Z radius:XY radius<br>mean lumen Z radius:XY radius<br>mean nuclear Z radius:XY radius<br>standard deviation nuclear Z radius:XY radius | 1. Divide the Z radius by the XY radius. |
| 3D radius | cyst 3D radius standard deviation<br>cyst 3D radius coefficient of variation | 1. Calculate the image slice where the object has the largest area.<br>2. Calculate the center of the object on this slice.<br>3. Measure the distance between all boundary points (on this slice) and the center.<br>4. Take the standard deviation or coefficient of variation. |
| external cell height ($\mu\text{m}$) | external cell height | 1. For every cyst coordiante, calculate the distance to the nearest lumen coordinate.<br>2. Take the mean. |
| external cell width ($\mu\text{m}$) | mean external cell width<br>standard deviation external cell width | 1. Calculate the center of each nucleus.<br>2. For each nucleus, calculate the distance to the nearest nucleus center. |
| major axis ($\mu\text{m}$) | cyst major axis<br>mean lumen major axis<br>mean nuclear major axis<br>standard deviation nuclear major axis | 1. Calculate the image slice where the object has the largest area.<br>2. Use MATLAB's regionprops function to calculate the major axis of the object on that slice.<br>3. Divide by 2.<br>4. Multiply by the size of the voxel in XY. |
| minor axis ($\mu\text{m}$) | cyst minor axis<br>mean lumen minor axis<br>mean nuclear minor axis<br>standard deviation nuclear minor axis | 1. Calculate the image slice where the object has the largest area.<br>2. Use MATLAB's regionprops function to calculate the minor axis of the object on that slice.<br>3. Divide by 2.<br>4. Multiply by the size of the voxel in XY. |
| major:minor axis | cyst major:minor axis<br>mean lumen major:minor axis<br>mean nuclear major:minor axis<br>standard deviation nuclear major:minor axis | 1. Divide the major axis by the minor axis. |
| solidity | cyst solidity<br>mean lumen solidity<br>mean nuclear solidity<br>standard deviation nuclear solidity | 1. Use MATLAB's regionprops function to calculate the solidity of the 3D object. |
| eccentricity | cyst eccentricity<br>mean lumen eccentricity<br>mean nuclear eccentricity<br>standard deviation nuclear eccentricity | 1. Calculate the image slice where the object has the largest area.<br>2. Use MATLAB's regionprops function to calculate the eccentricity of the object on that slice. |
| number | number of lumens<br>number of nuclei<br>number of internal nuclei<br>number of external nuclei | 1. Count the number of objects. |
| density | density of lumens<br>density of nuclei | 1. Divide the number of objects by the cyst volume. |

**Supplemental Table 2:** Perturbation Drugs and Concentrations.

| Drug | Concentrations |
| --- | --- |
| selinexor | 0.1, 0.3 μM |
| lapatinib | 40, 130, 400 nM |
| givinostat | 0.1 μM |
| PI-103 | 0.1, 0.3 μM |
| idelalisib | 3, 10, 30 μM |
| mardepodect | 0.1, 1.0 μM |
| sumatriptan succinate | 0.1, 0.3, 1.0 μM |
| Aurora A Inhibitor I | 0.1, 0.3, 1.0 μM |
| orantinib | 0.5, 1.6, 5.0 μM |
| Y-27632 | 10 μM |
| NSC23766 | 30 μM |
| blebbistatin | 10 μM |
| HGF | 5, 20 ng/mL |

**Supplemental Table 3:**
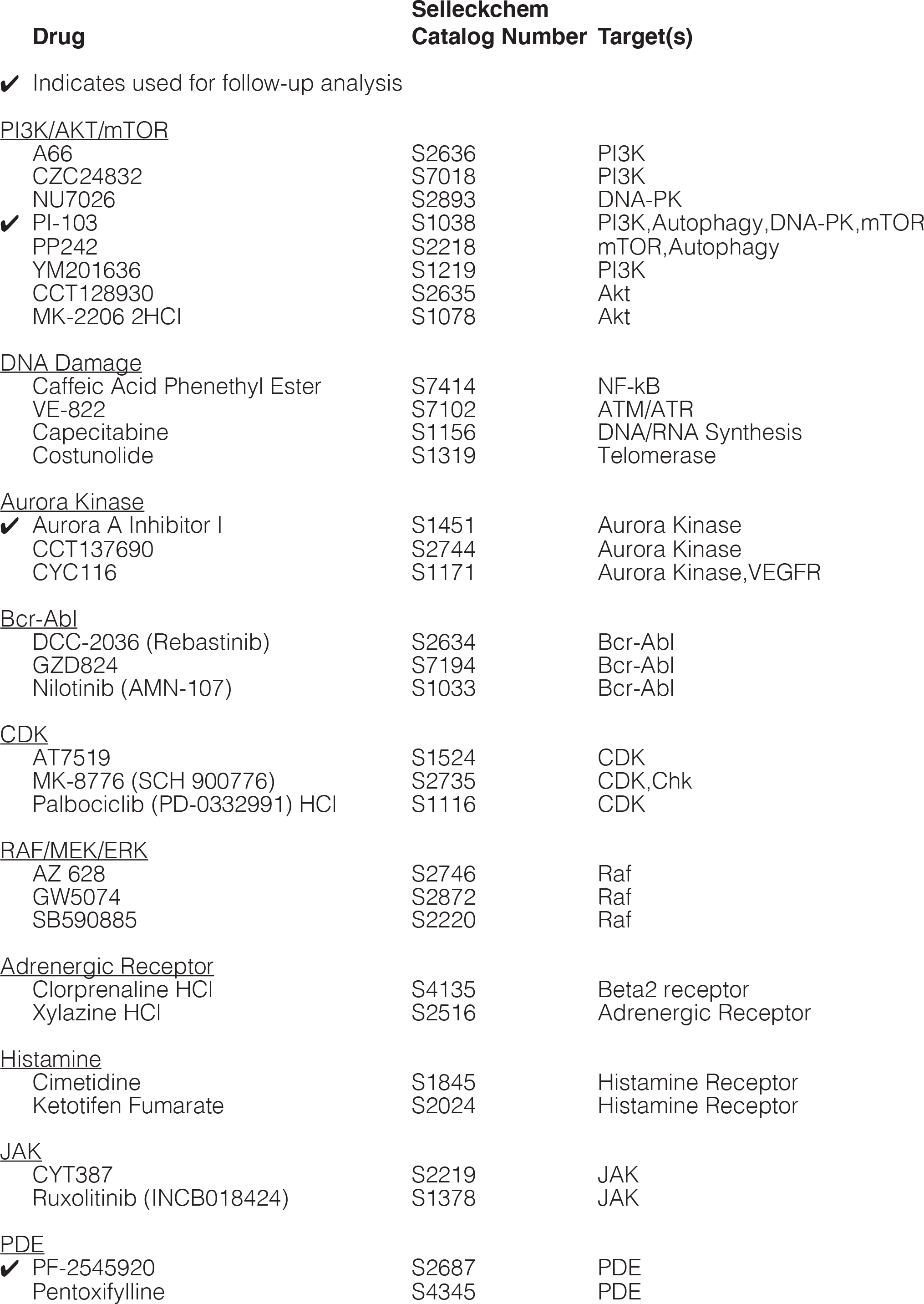
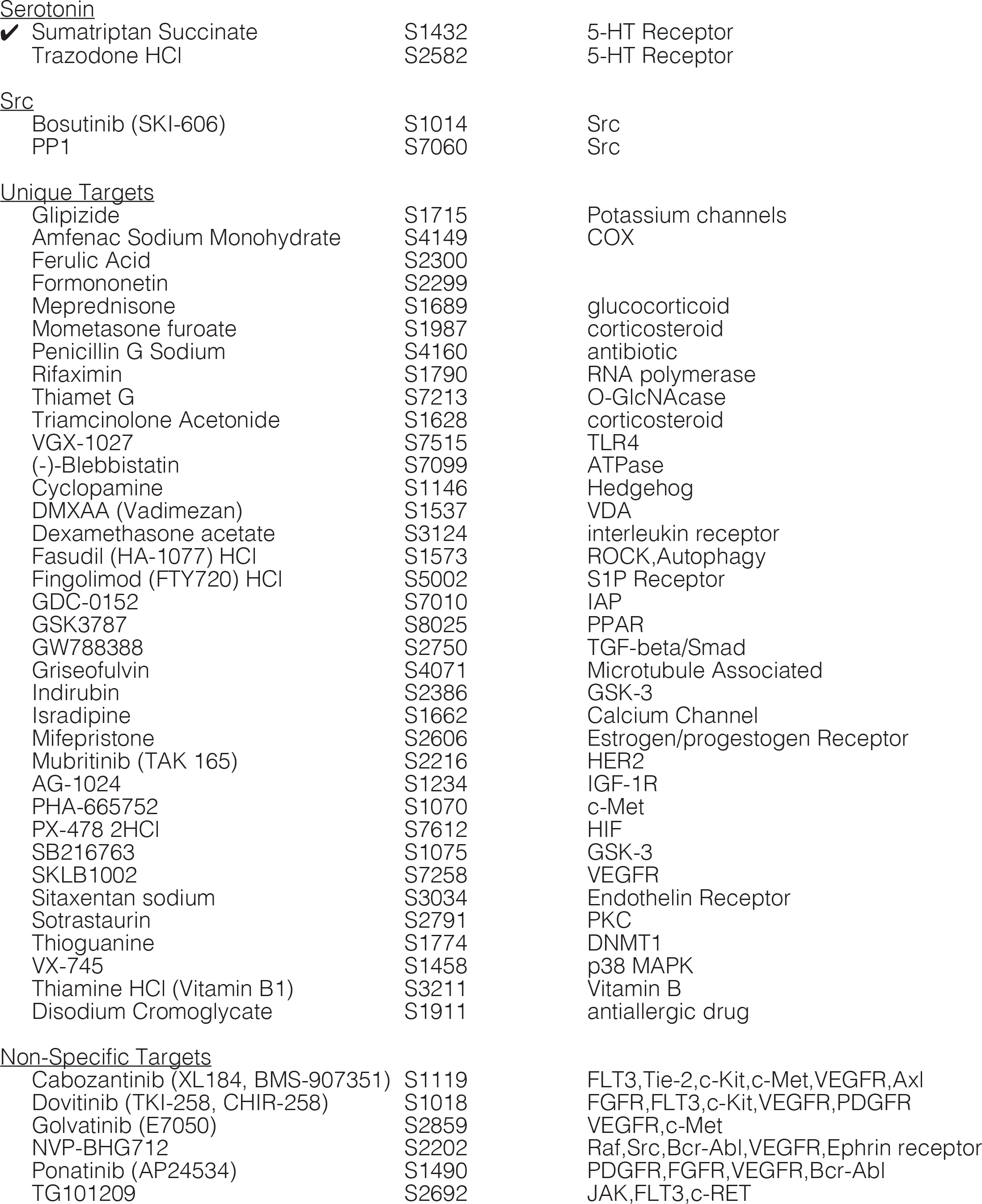
Hits for Larger MDCK Cysts from Drug Screen, Grouped by Target.

| Drug | Selleckchem<br>Catalog Number | Target(s) |
| --- | --- | --- |
| ✓ Indicates used for follow-up analysis |  |  |
| <u>PI3K/AKT/mTOR</u> |  |  |
| A66 | S2636 | PI3K |
| CZC24832 | S7018 | PI3K |
| NU7026 | S2893 | DNA-PK |
| ✓ PI-103 | S1038 | PI3K, Autophagy, DNA-PK, mTOR |
| PP242 | S2218 | mTOR, Autophagy |
| YM201636 | S1219 | PI3K |
| CCT128930 | S2635 | Akt |
| MK-2206 2HCl | S1078 | Akt |
| <u>DNA Damage</u> |  |  |
| Caffeic Acid Phenethyl Ester | S7414 | NF-kB |
| VE-822 | S7102 | ATM/ATR |
| Capecitabine | S1156 | DNA/RNA Synthesis |
| Costunolide | S1319 | Telomerase |
| <u>Aurora Kinase</u> |  |  |
| ✓ Aurora A Inhibitor I | S1451 | Aurora Kinase |
| CCT137690 | S2744 | Aurora Kinase |
| CYC116 | S1171 | Aurora Kinase, VEGFR |
| <u>Bcr-Abl</u> |  |  |
| DCC-2036 (Rebastinib) | S2634 | Bcr-Abl |
| GZD824 | S7194 | Bcr-Abl |
| Nilotinib (AMN-107) | S1033 | Bcr-Abl |
| <u>CDK</u> |  |  |
| AT7519 | S1524 | CDK |
| MK-8776 (SCH 900776) | S2735 | CDK, Chk |
| Palbociclib (PD-0332991) HCl | S1116 | CDK |
| <u>RAF/MEK/ERK</u> |  |  |
| AZ 628 | S2746 | Raf |
| GW5074 | S2872 | Raf |
| SB590885 | S2220 | Raf |
| <u>Adrenergic Receptor</u> |  |  |
| Clorprenaline HCl | S4135 | Beta2 receptor |
| Xylazine HCl | S2516 | Adrenergic Receptor |
| <u>Histamine</u> |  |  |
| Cimetidine | S1845 | Histamine Receptor |
| Ketotifen Fumarate | S2024 | Histamine Receptor |
| <u>JAK</u> |  |  |
| CYT387 | S2219 | JAK |
| Ruxolitinib (INCB018424) | S1378 | JAK |
| <u>PDE</u> |  |  |
| ✓ PF-2545920 | S2687 | PDE |
| Pentoxifylline | S4345 | PDE |
| Serotonin |  |  |
| ✓ Sumatriptan Succinate | S1432 | 5-HT Receptor |
| Trazodone HCl | S2582 | 5-HT Receptor |
| Src |  |  |
| Bosutinib (SKI-606) | S1014 | Src |
| PP1 | S7060 | Src |
| Unique Targets |  |  |
| Glipizide | S1715 | Potassium channels |
| Amfenac Sodium Monohydrate | S4149 | COX |
| Ferulic Acid | S2300 |  |
| Formononetin | S2299 |  |
| Meprednisone | S1689 | glucocorticoid |
| Mometasone furoate | S1987 | corticosteroid |
| Penicillin G Sodium | S4160 | antibiotic |
| Rifaximin | S1790 | RNA polymerase |
| Thiamet G | S7213 | O-GlcNAcase |
| Triamcinolone Acetonide | S1628 | corticosteroid |
| VGX-1027 | S7515 | TLR4 |
| (-)-Blebbistatin | S7099 | ATPase |
| Cyclopamine | S1146 | Hedgehog |
| DMXAA (Vadimezan) | S1537 | VDA |
| Dexamethasone acetate | S3124 | interleukin receptor |
| Fasudil (HA-1077) HCl | S1573 | ROCK, Autophagy |
| Fingolimod (FTY720) HCl | S5002 | S1P Receptor |
| GDC-0152 | S7010 | IAP |
| GSK3787 | S8025 | PPAR |
| GW788388 | S2750 | TGF-beta/Smad |
| Griseofulvin | S4071 | Microtubule Associated |
| Indirubin | S2386 | GSK-3 |
| Isradipine | S1662 | Calcium Channel |
| Mifepristone | S2606 | Estrogen/progestogen Receptor |
| Mubritinib (TAK 165) | S2216 | HER2 |
| AG-1024 | S1234 | IGF-1R |
| PHA-665752 | S1070 | c-Met |
| PX-478 2HCl | S7612 | HIF |
| SB216763 | S1075 | GSK-3 |
| SKLB1002 | S7258 | VEGFR |
| Sitaxentan sodium | S3034 | Endothelin Receptor |
| Sotrastaurin | S2791 | PKC |
| Thioguanine | S1774 | DNMT1 |
| VX-745 | S1458 | p38 MAPK |
| Thiamine HCl (Vitamin B1) | S3211 | Vitamin B |
| Disodium Cromoglycate | S1911 | antiallergic drug |

|  |  |  |
| --- | --- | --- |
| Cabozantinib (XL184, BMS-907351) | S1119 | FLT3, Tie-2, c-Kit, c-Met, VEGFR, Axl |
| Dovitinib (TKI-258, CHIR-258) | S1018 | FGFR, FLT3, c-Kit, VEGFR, PDGFR |
| Golvatinib (E7050) | S2859 | VEGFR, c-Met |
| NVP-BHG712 | S2202 | Raf, Src, Bcr-Abl, VEGFR, Ephrin receptor |
| Ponatinib (AP24534) | S1490 | PDGFR, FGFR, VEGFR, Bcr-Abl |
| TG101209 | S2692 | JAK, FLT3, c-RET |

**Supplemental Table 4:**
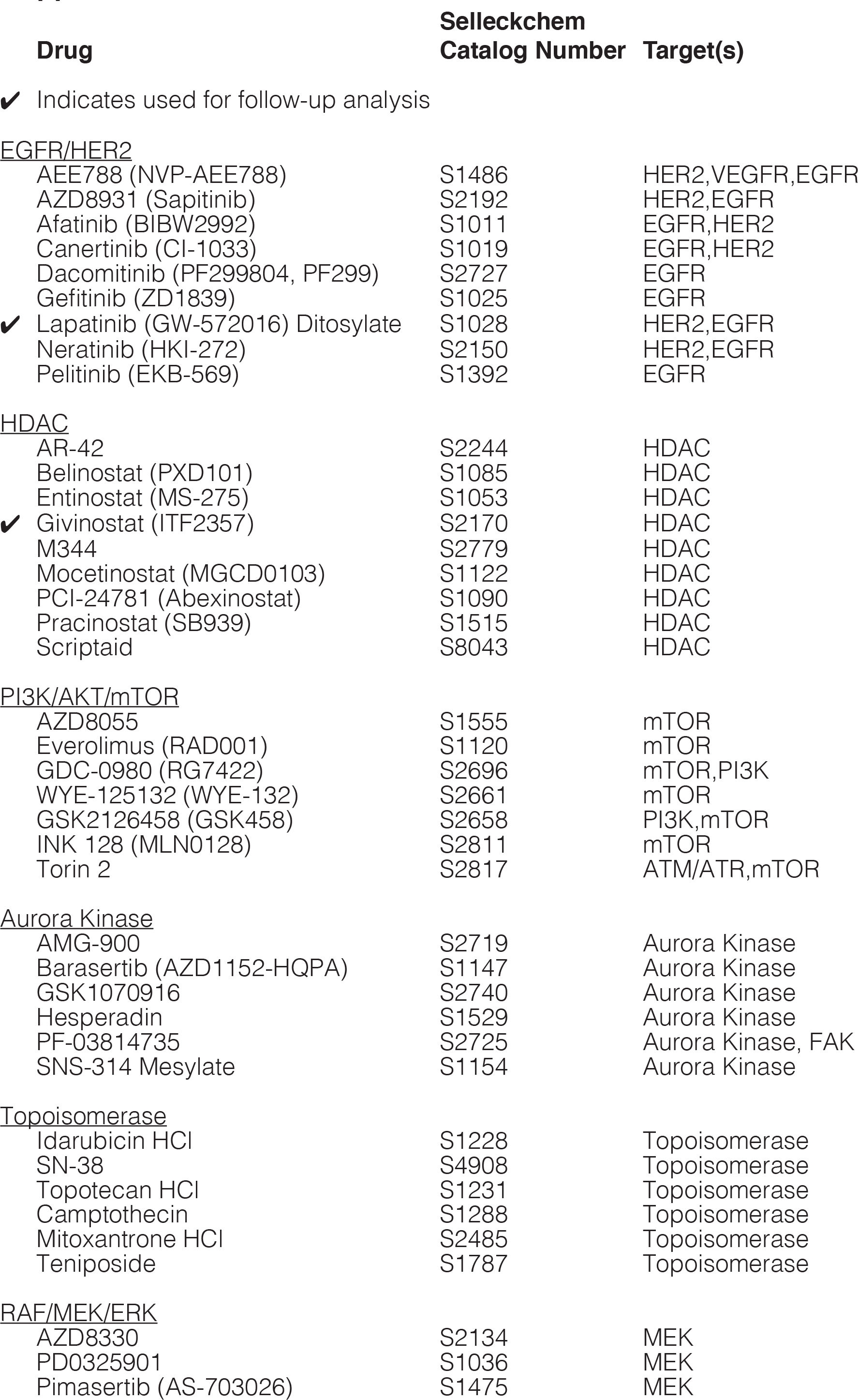
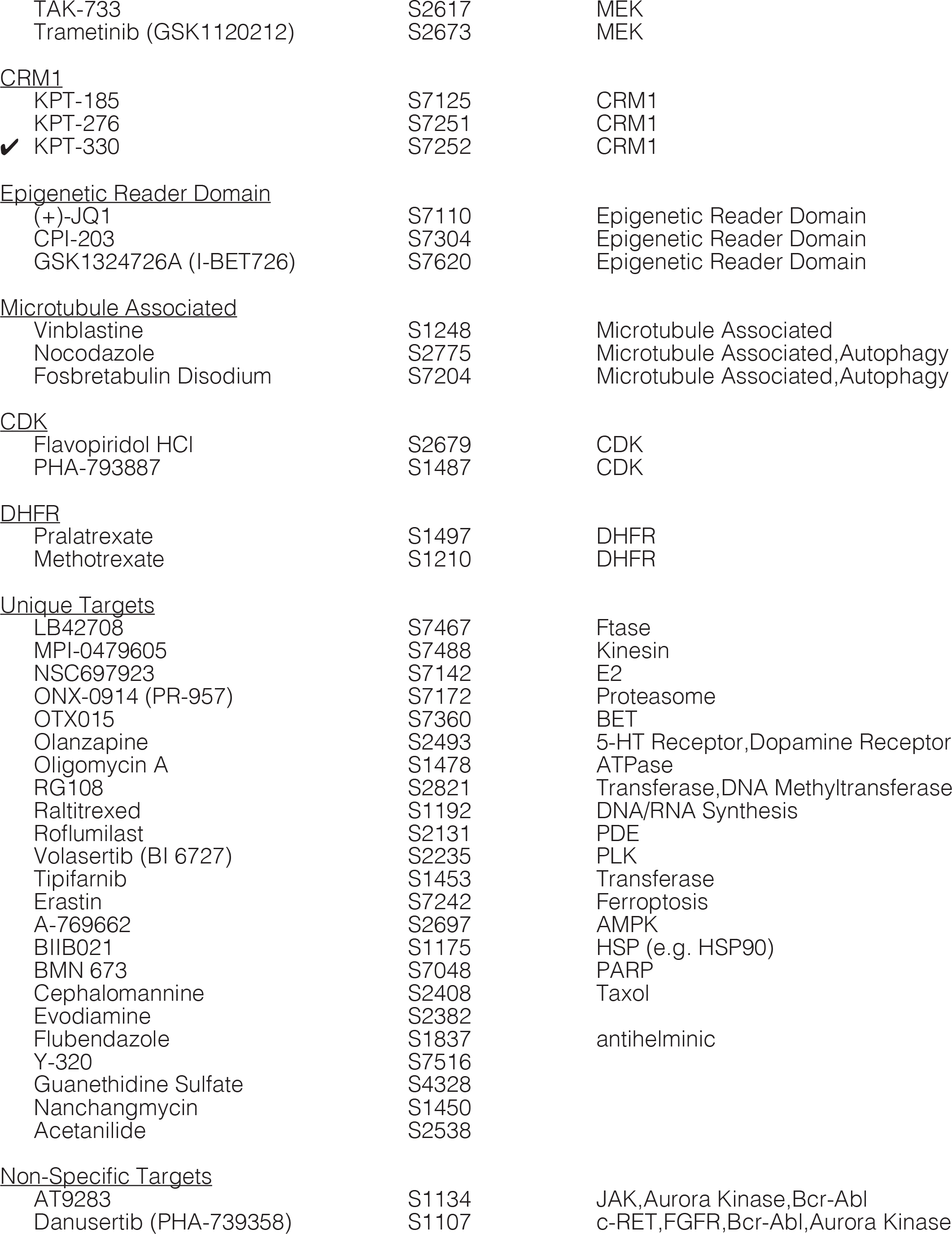
Hits for Smaller MDCK Cysts from Drug Screen, Grouped by Target.

**Supplemental Figure 1:**
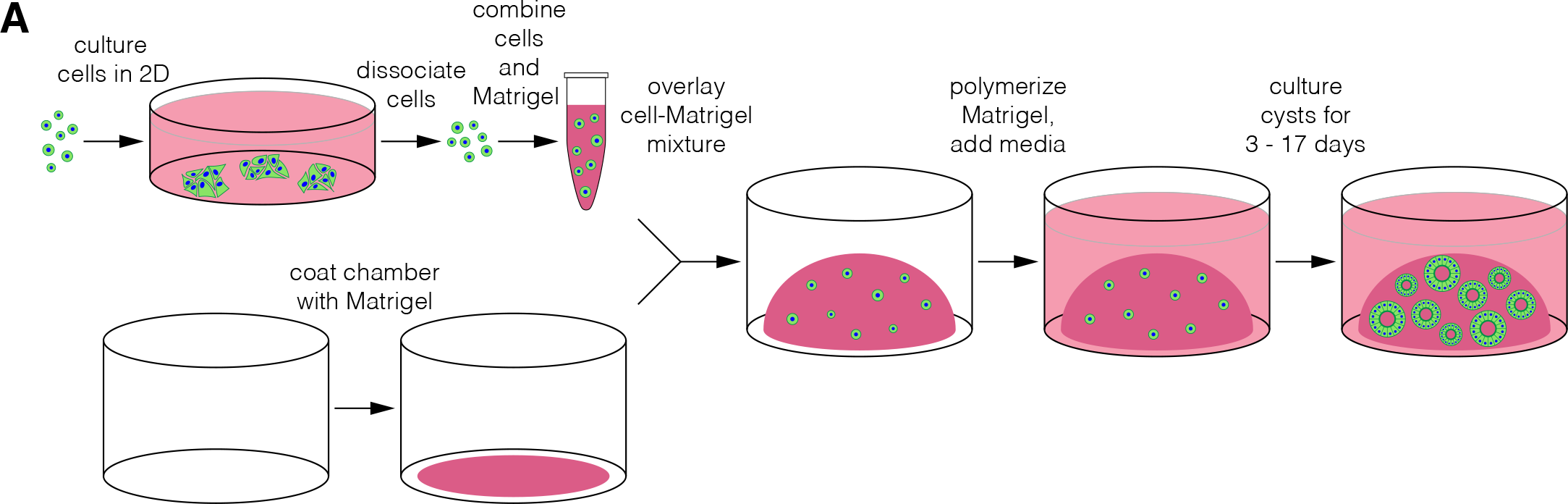
Schematic for MDCK cyst culture technique. **A.** MOCK cells are maintained in two-dimensional culture. When the cells are sufficiently confluent, they are dissociated into a single cell suspension. Cells are added to liquid Matrigel and the cell-Matrigel mixture is plated into a cell culture chamber already coated with pure Matrigel. After the Matrigel has polymerized, media can be added and the cysts can be cultured for at least 17 days. See methods for more information.

**Supplemental Figure 2:**
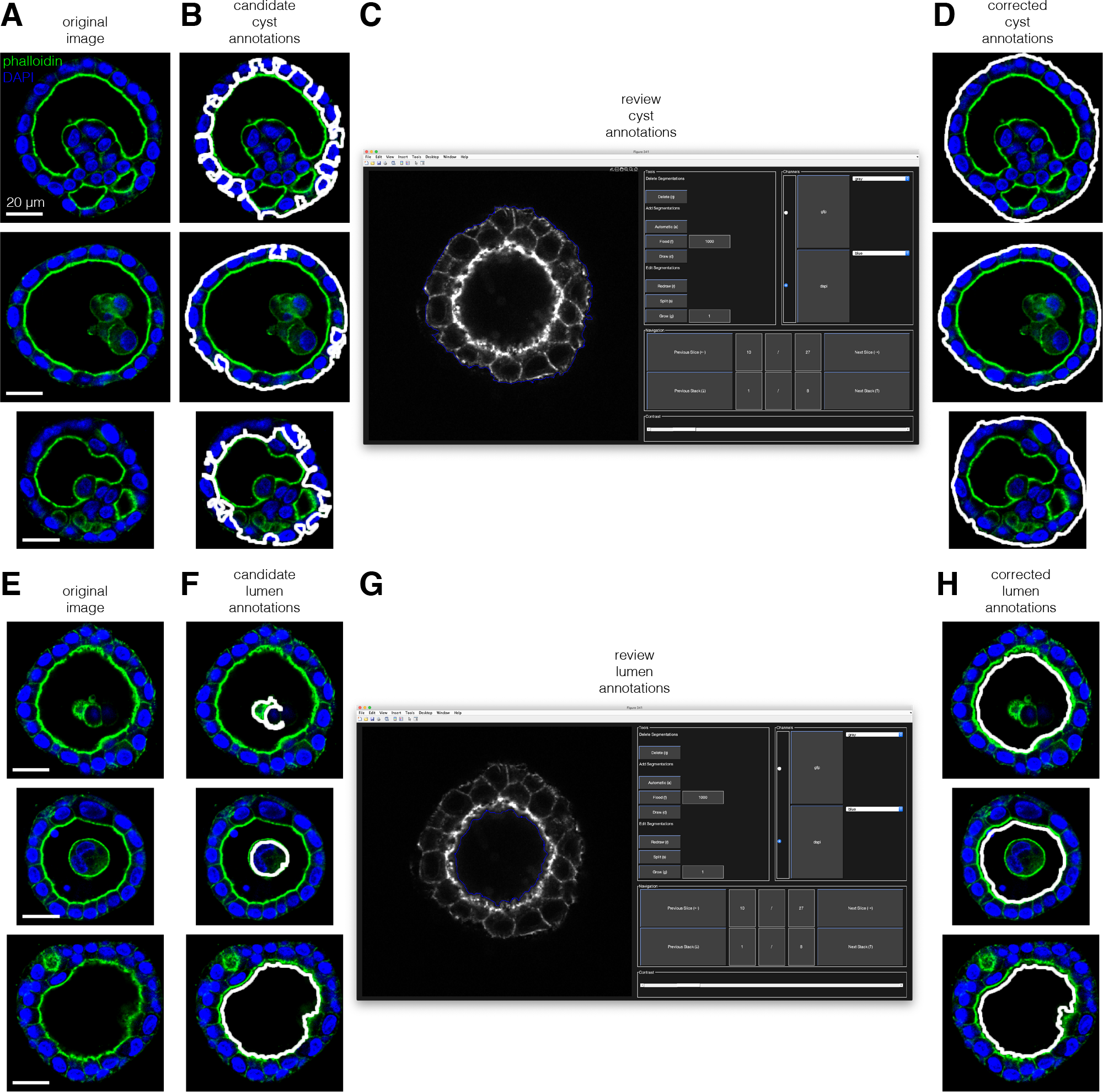
Manually correcting 2D cyst and lumen annotations. **A.** Example images of MOCK cysts. **B.** Example candidate annotations for the cyst boundary. **C.** Our user interface for viewing and correcting 2O annotations. **D.** Example annotations for the cyst boundary after correction. **E.** Example images of MOCK cysts. **F.** Example candidate annotations for the lumen boundaries. **G.** Our user interface for viewing and correcting 2O annotations. **H.** Example annotations for the lumen boundaries after correction.

**Supplemental Figure 3:**
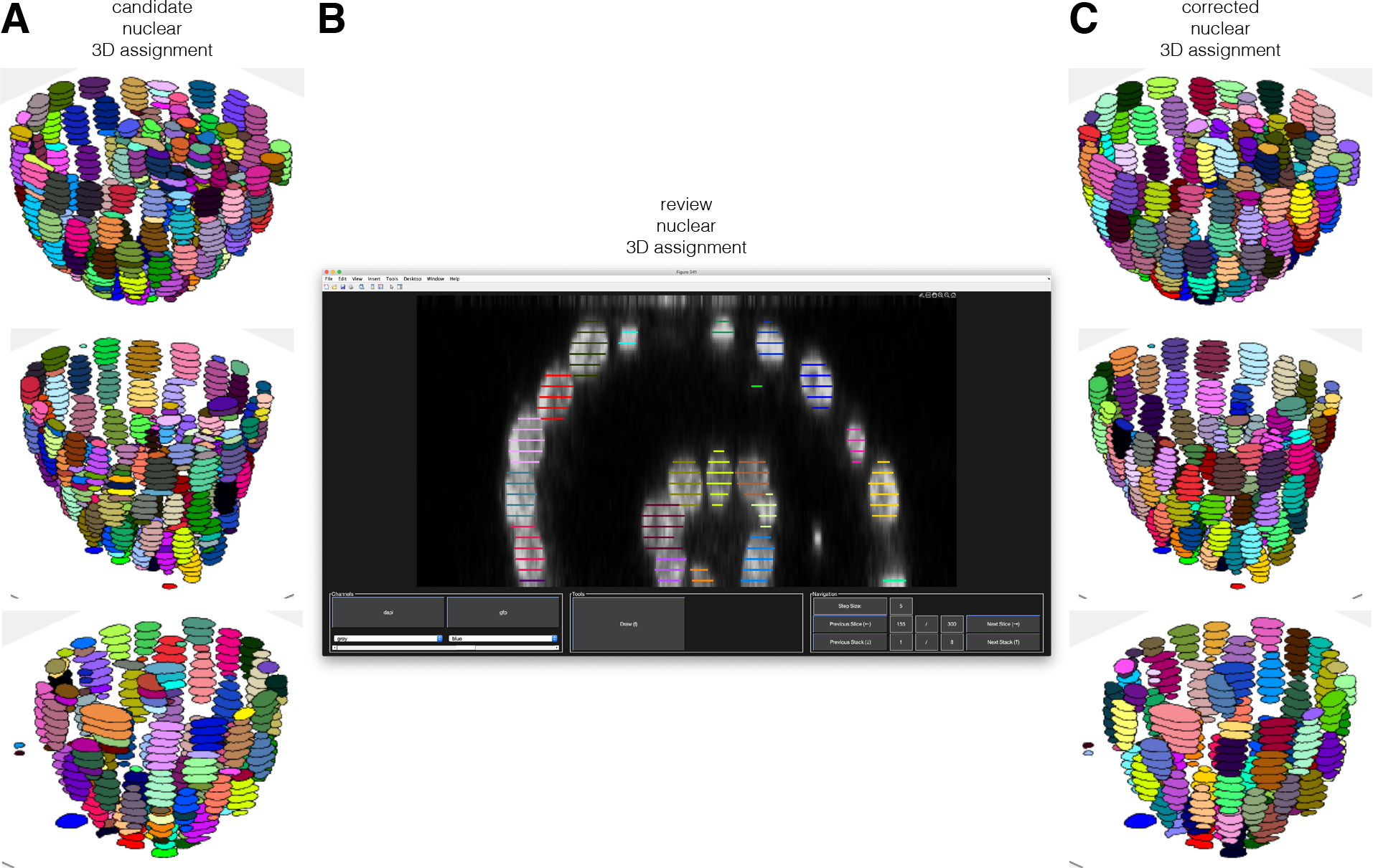
Manually correcting 3D nuclear annotations. **A.** Example candidate nuclear annotations, color-coded by which 3O object they belong to. **B.** Our user interface for viewing and correcting 3O nuclear annotations. **C.** Example nuclear annotations after they have been manually corrected.

**Supplemental Figure 4:**
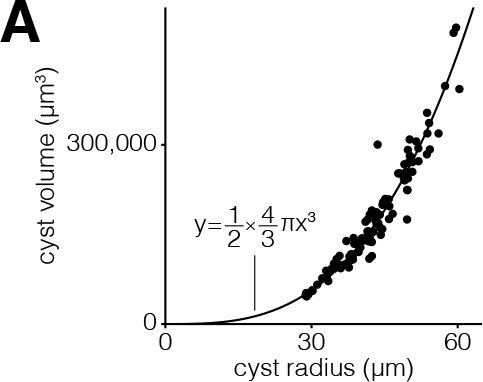
Cyst volume scales with cyst radius to the third. **A.** Cyst volume versus cyst radius for 7-11 day old MOCK cysts. Reference line indicates the relationship between radius and volume of half a sphere.

**Supplemental Figure 5:**
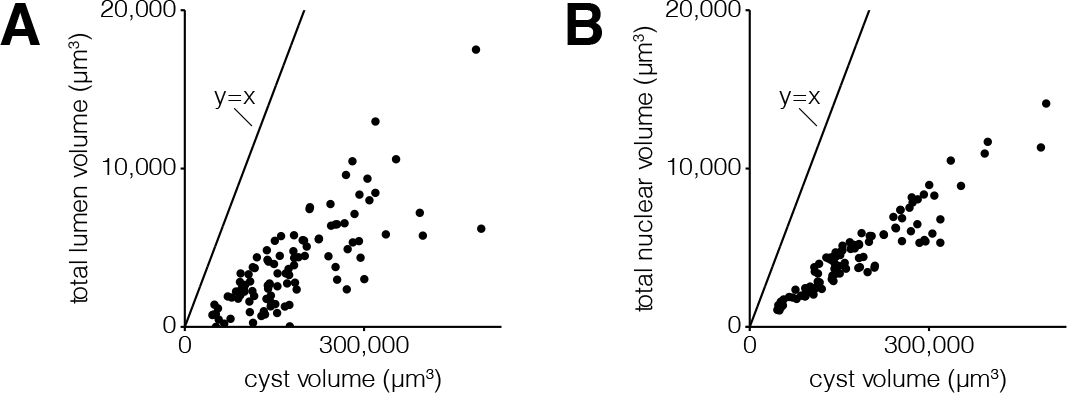
Total lumen and nuclear volumes are less than cyst volume. **A.** Total lumen volume versus cyst volume for 7-11 day old MOCK cysts. Reference line indicates y = x. **B.** Total nuclear volume versus cyst volume for 7-11 day old MOCK cysts. Reference line indicates y = x.

**Supplemental Figure 6:**
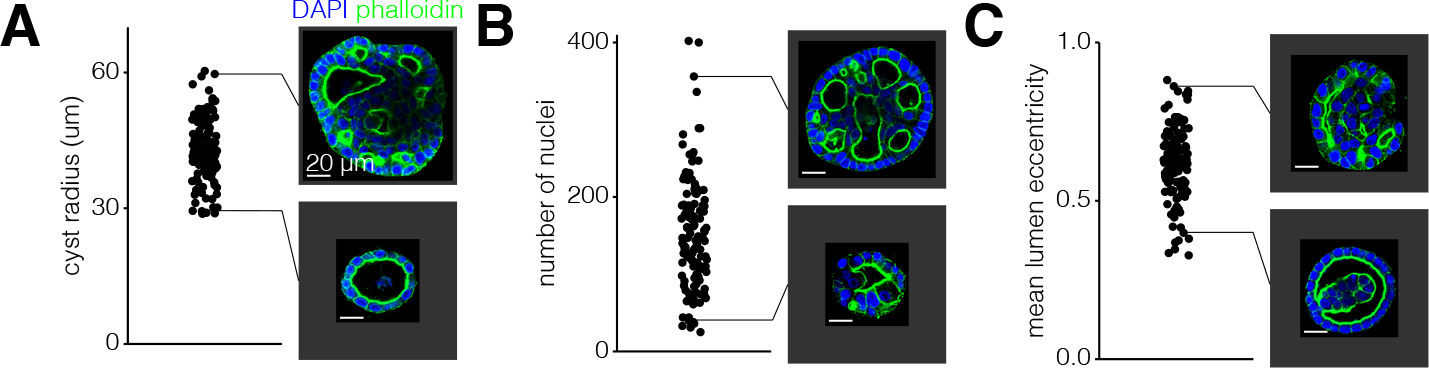
Examples MDCK cysts with high and low values for cyst morphological features. **A.** Cyst radius for 7-11 day old MOCK cysts with example images. **B.** Number of nuclei for 7-11 day old MOCK cysts with example images. **C.** Mean lumen eccentricity for 7-11 day old MOCK cysts with example images.

**Supplemental Figure 7:**
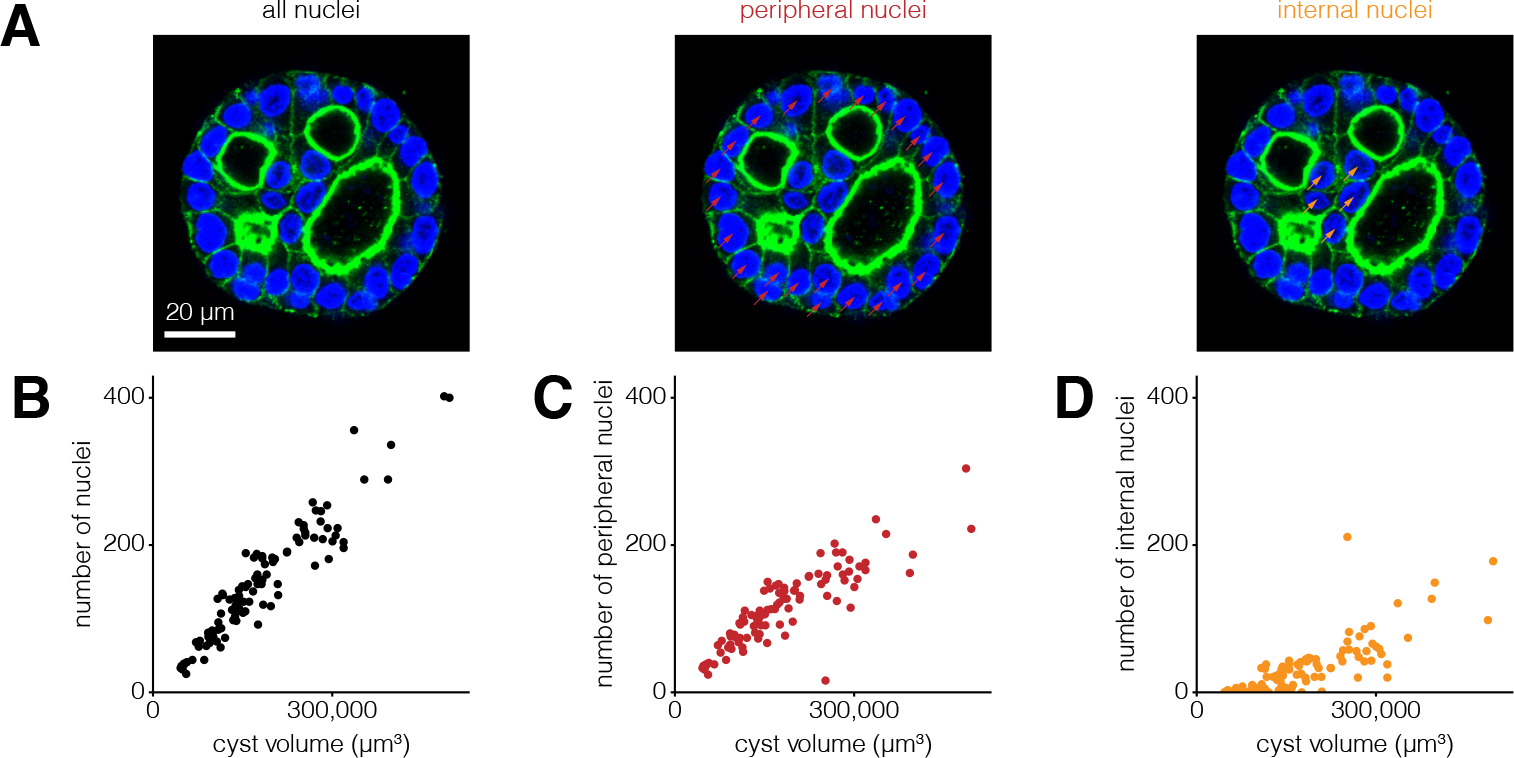
Peripheral scale sublinearly and internal nuclei scale superlinearly with cyst volume. **A.** Example MOCK cysts with peripheral nuclei annotated with a red dot and internal nuclei annotated with an orange dot. **B.** Number of nuclei versus cyst volume for 7-11 day old MOCK cysts. **C.** Number of peripheral nuclei versus cyst volume for 7-11 day old MOCK cysts. **D.** Number of internal nuclei versus cyst volume for 7-11 day old MOCK cysts.

**Supplemental Figure 8:**
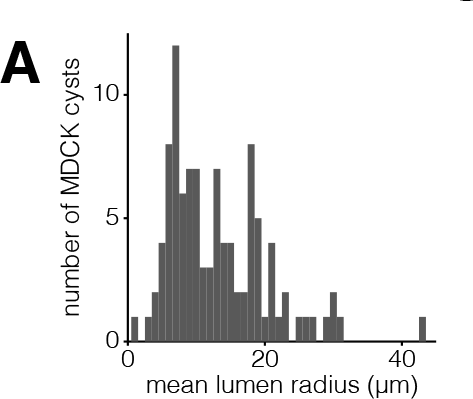
MDCK cysts may have a minimum size for lumens. **A.** Histogram (with a bin width of 1 um) of mean lumen radius for MOCK cysts cultured for 7-11 days.

**Supplemental Figure 9:**
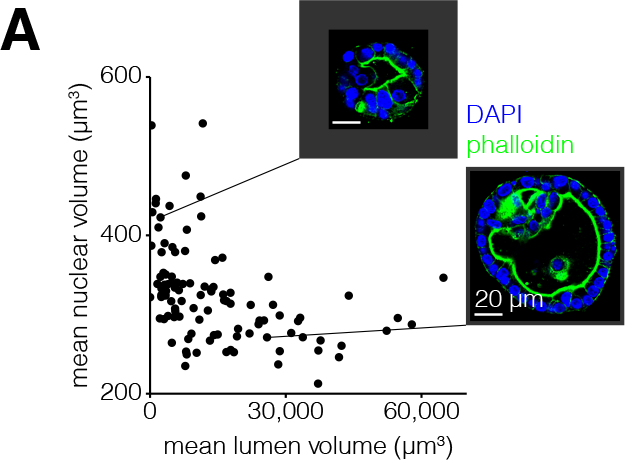
Mean nuclear size is inversely correlated with mean lumen size. **A.** Mean nuclear volume versus mean lumen volume for 7-11 day old MOCK cysts with example images.

**Supplemental Figure 10:**
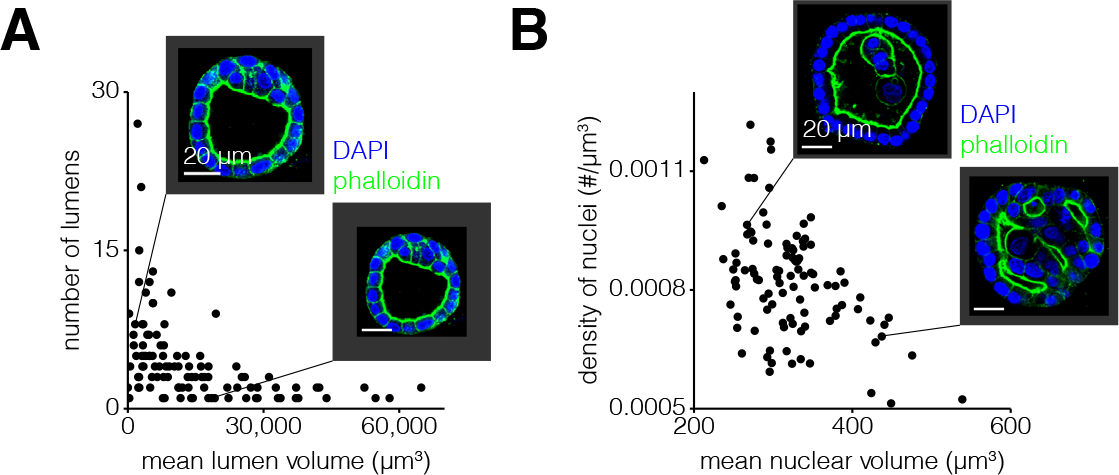
The number of lumens and nuclei are inversely correlated with their mean volume. **A.** Number of lumens versus mean lumen volume for 7-11 day old MOCK cysts with example images. **B.** Oensity of nuclei versus mean nuclear volume for 7-11 day old MOCK cysts with example images.

**Supplemental Figure 11:**
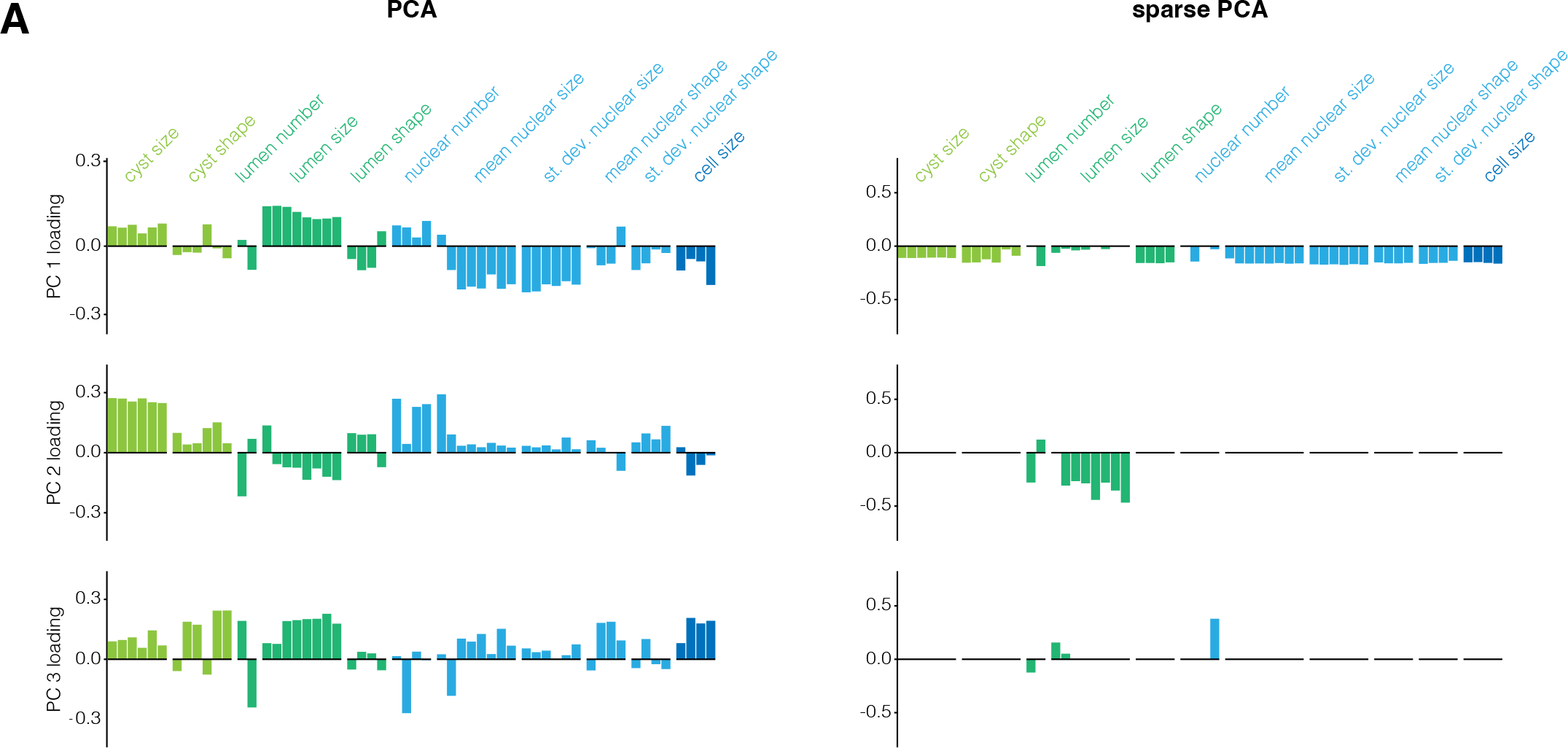
Sparse PCA reveals similar constraints to traditional PCA. **A.** loading of each feature on principal components one through three for both standard PCA and sparse PCA. Each feature is color-coded by what structure (cyst, lumen, nucleus, or cell) it describes.

**Supplemental Figure 12:**
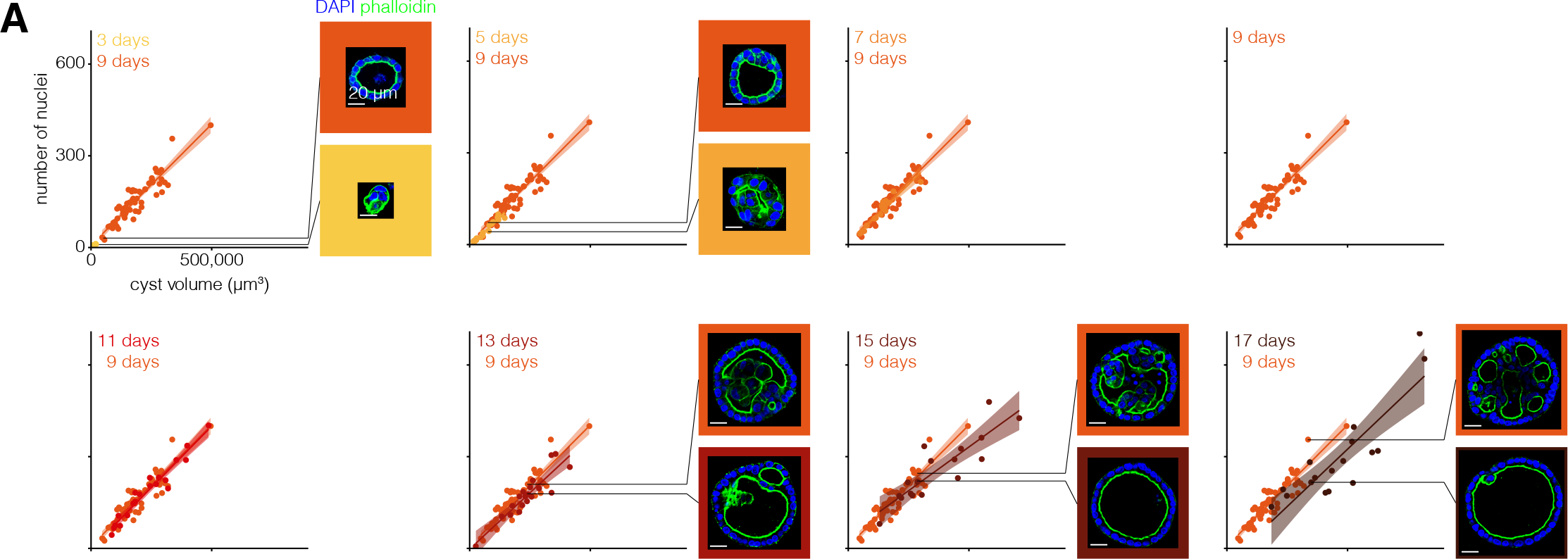
Constraint on number of nuclei and cyst volume varies with MDCK cyst age. **A.** Number of nuclei versus cyst volume for MOCK cysts of each age. Each age is represented by one color, and 9 day old MOCK cysts are repeated on each graph for reference. The line represents the line of best fit and the shaded area represents the 95% confidence interval. Example MOCK cysts of different ages with approximately the same volume and different numbers of nuclei are shown.

**Supplemental Figure 13:**
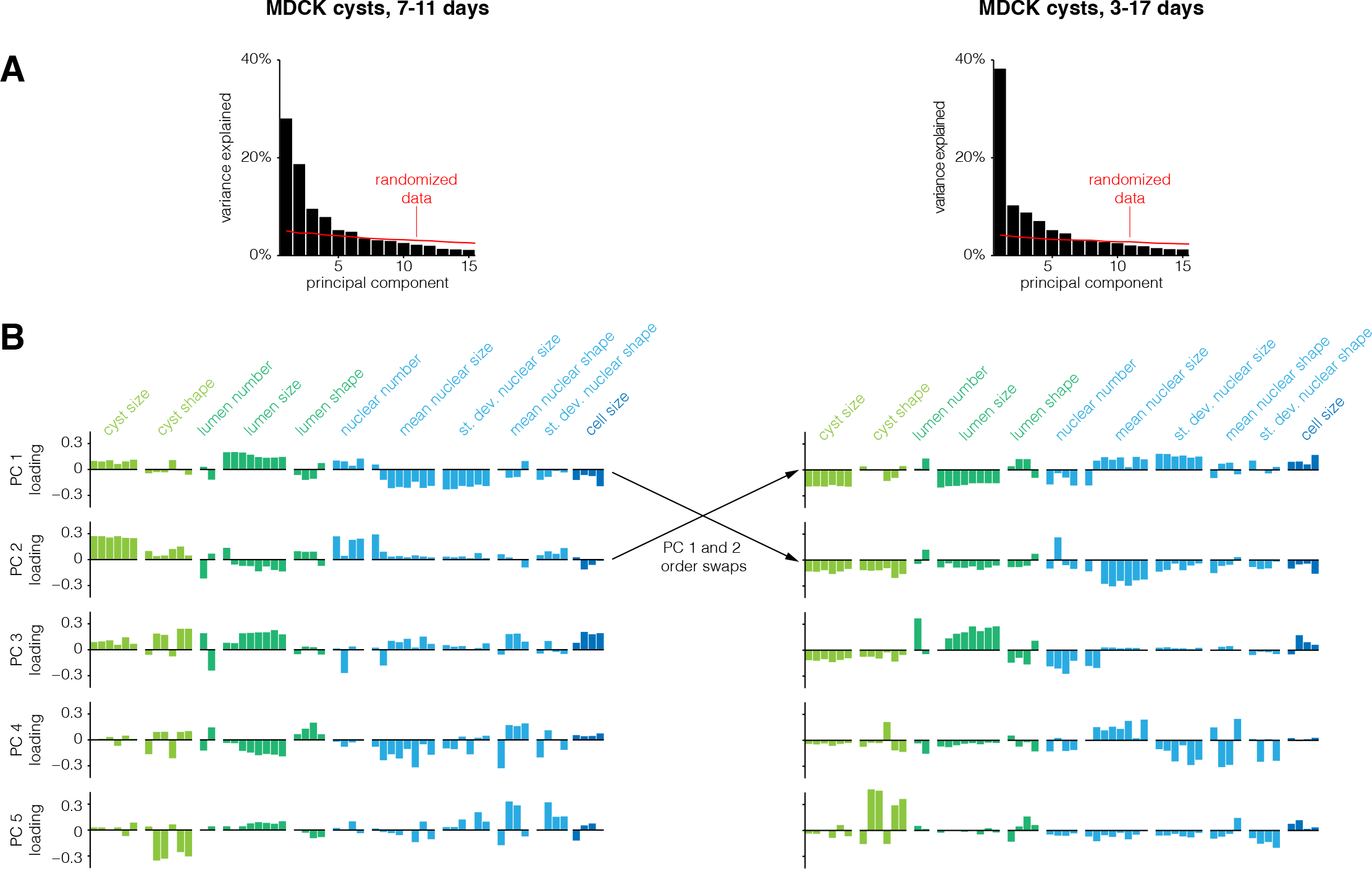
Constraints on MDCK cysts are consistent across cysts of all ages. **A.** Variance explained by each principal component. The red line indicates how much variance is explained when the data is randomized before PCA (see methods for details). **B.** loading of each feature on principal components one through five. Each feature is color-coded by what structure (cyst, lumen, nucleus, or cell) it describes.

**Supplemental Figure 14:**
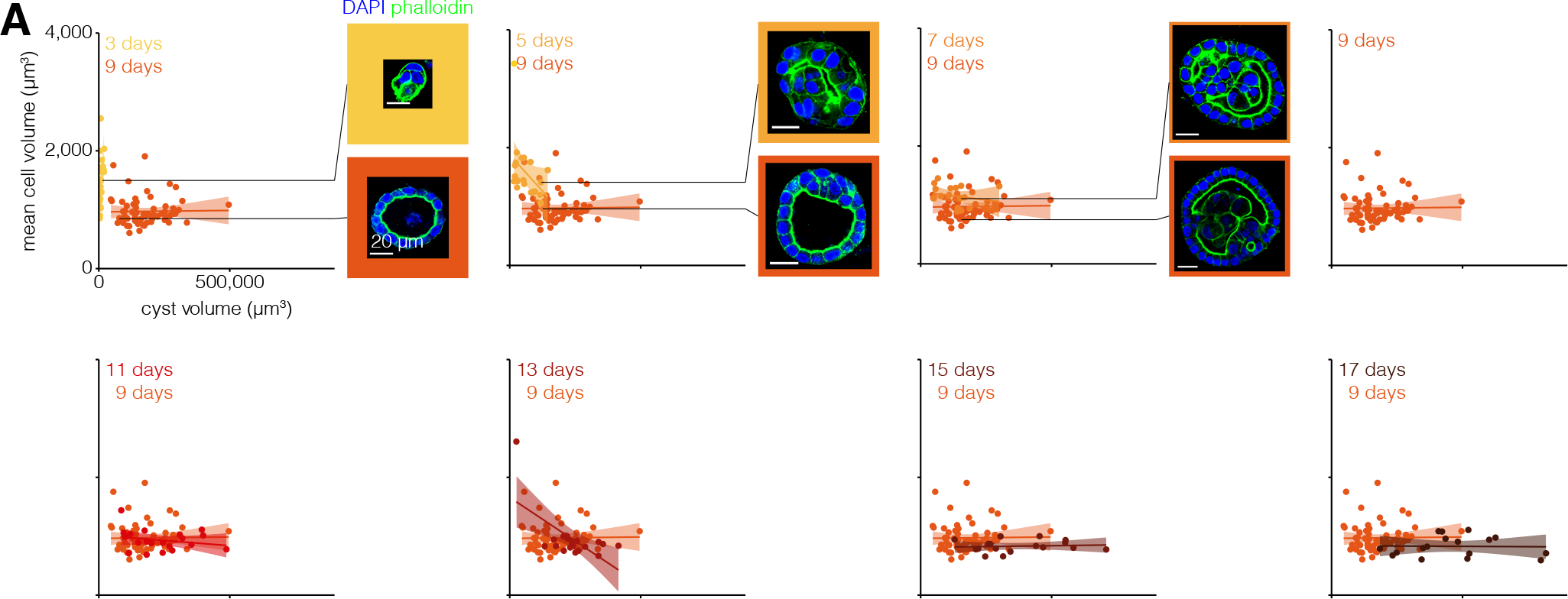
Constraint on cell volume and cyst volume varies with MDCK cyst age. **A.** Mean cell volume versus cyst volume for MOCK cysts of each age. Each age is represented by one color, and 9 day old MOCK cysts are repeated on each graph for reference. The line represents the line of best fit and the shaded area represents the 95% confidence interval. Example MOCK cysts of different ages with approximately the same cyst volume and different mean cell volume are shown.

**Supplemental Figure 15:**
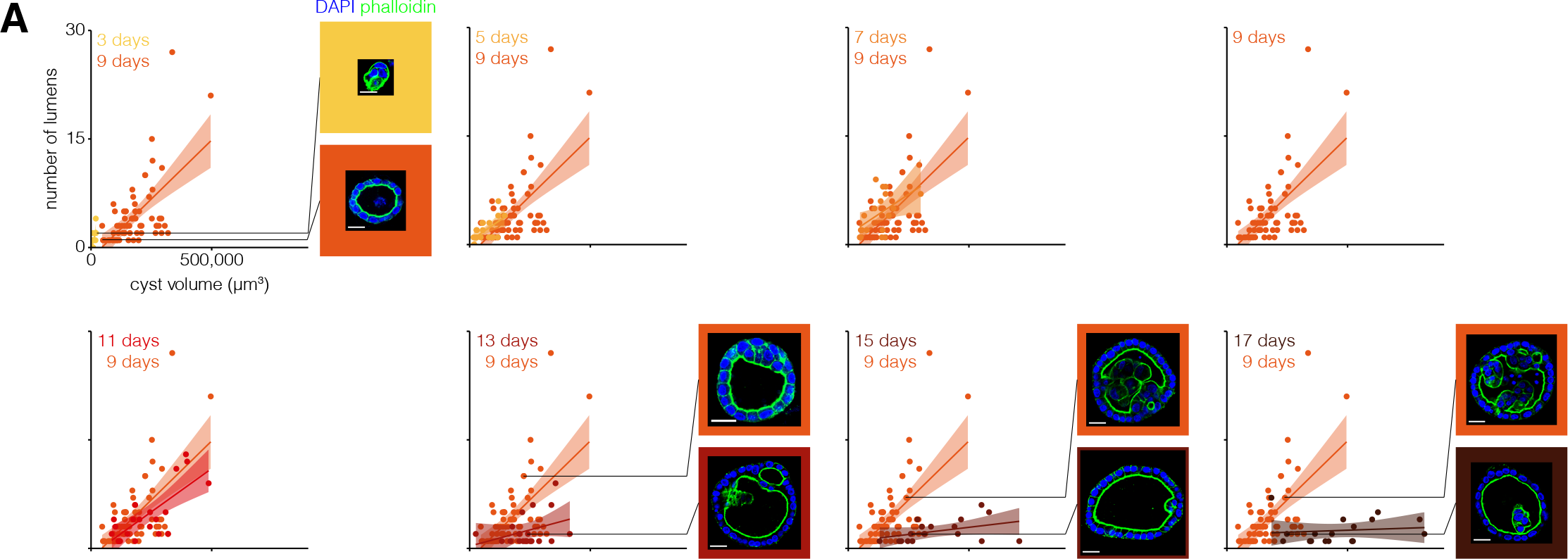
Constraint on number of lumens and cyst volume varies with MDCK cyst age. **A.** Number of lumens versus cyst volume for MOCK cysts of each age. Each age is represented by one color, and 9 day old MOCK cysts are repeated on each graph for reference. The line represents the line of best fit and the shaded area represents the 95% confidence interval. Example MOCK cysts of different ages with approximately the same cyst volume and different numbers of lumens are shown.

**Supplemental Figure 16:**
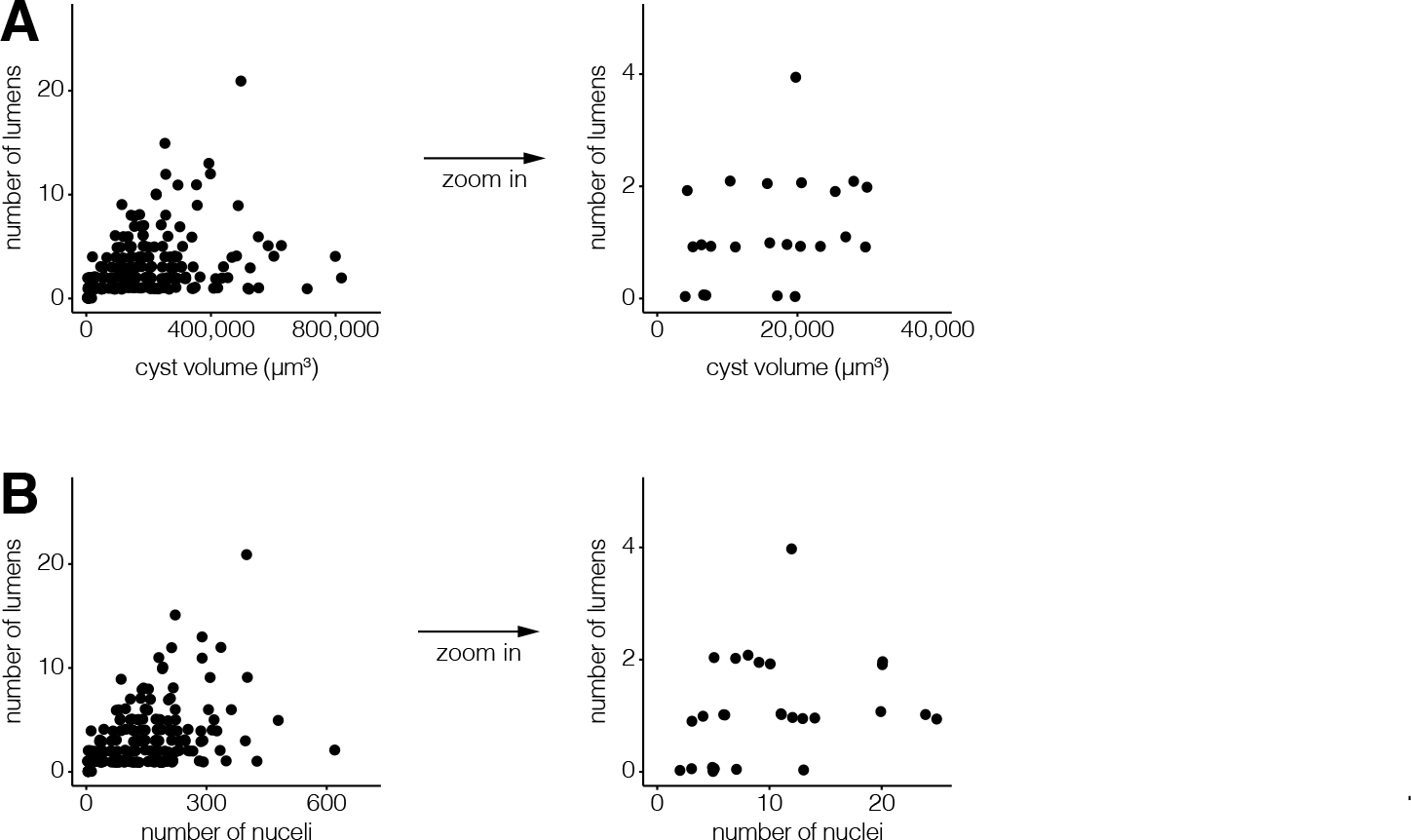
MDCK cysts develop a lumen when they have 7 cells and are 10,000 µm^3^. **A.** Number of lumens versus cyst volume for MOCK cysts with 3-17 days of growth. **B.** Number of lumens versus number of cells for MOCK cysts with 3-17 days of growth.

**Supplemental Figure 17:**
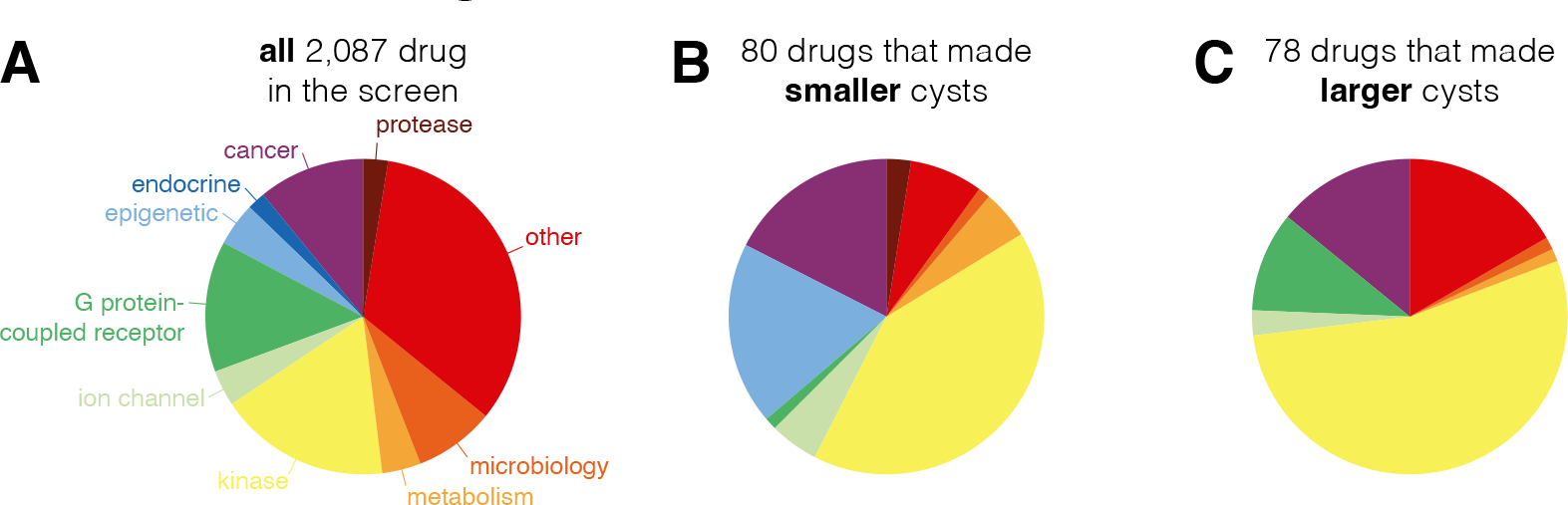
Proportions of drug categories for drug screen. **A.** Proportion of drugs categories for all drugs screened. **B.** Proportion of drug categories amongst drugs found to decrease cyst area. **C.** Proportion of drug categories amongst drugs found to increase cyst area.

**Supplemental Figure 18:**
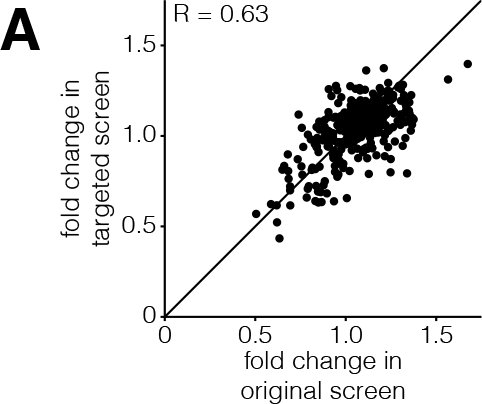
Correlation between original and targeted drug screen. **A.** For drugs screened in replicate (1/7th of all drugs screen), the fold change in the targeted screen versus the fold change in the original screen.

**Supplemental Figure 19:**
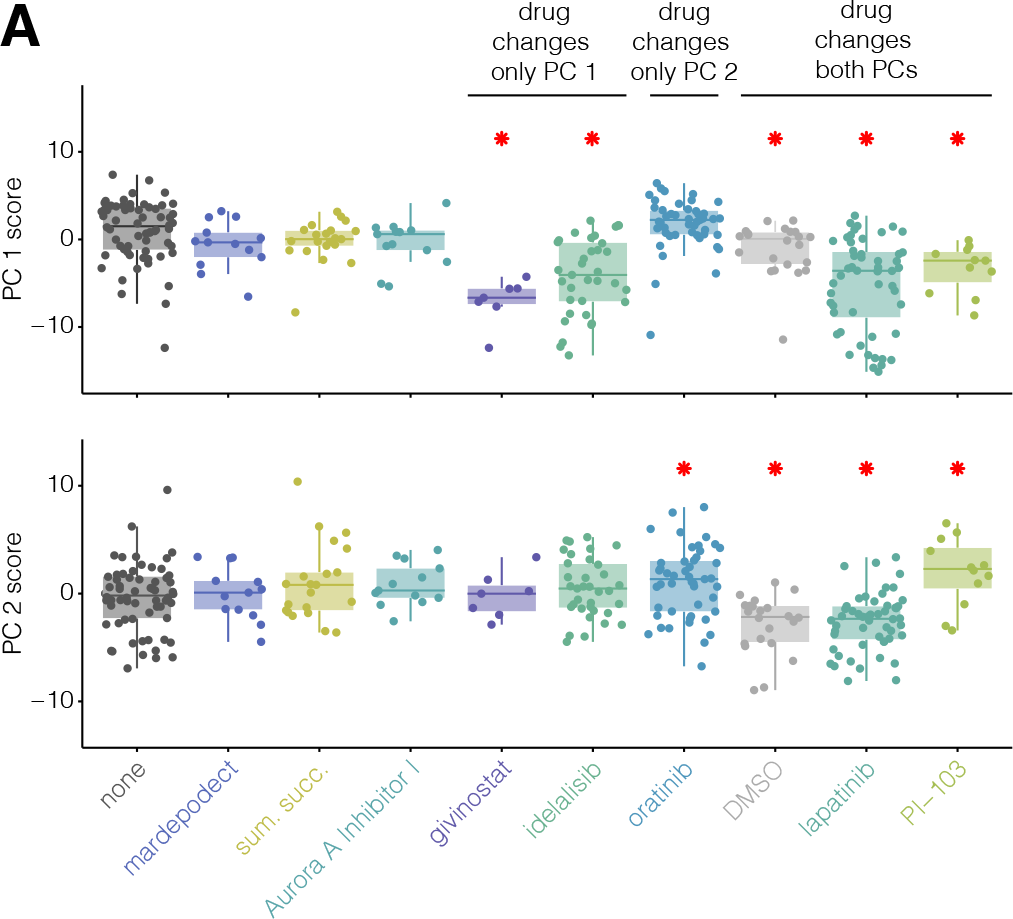
Perturbations can shift MDCK cysts along more than one principal component. **A.** Scores of MOCK cysts (of various perturbations) when projected into PC space calculated using unperturbed MOCK cysts.

**Supplemental Figure 20:**
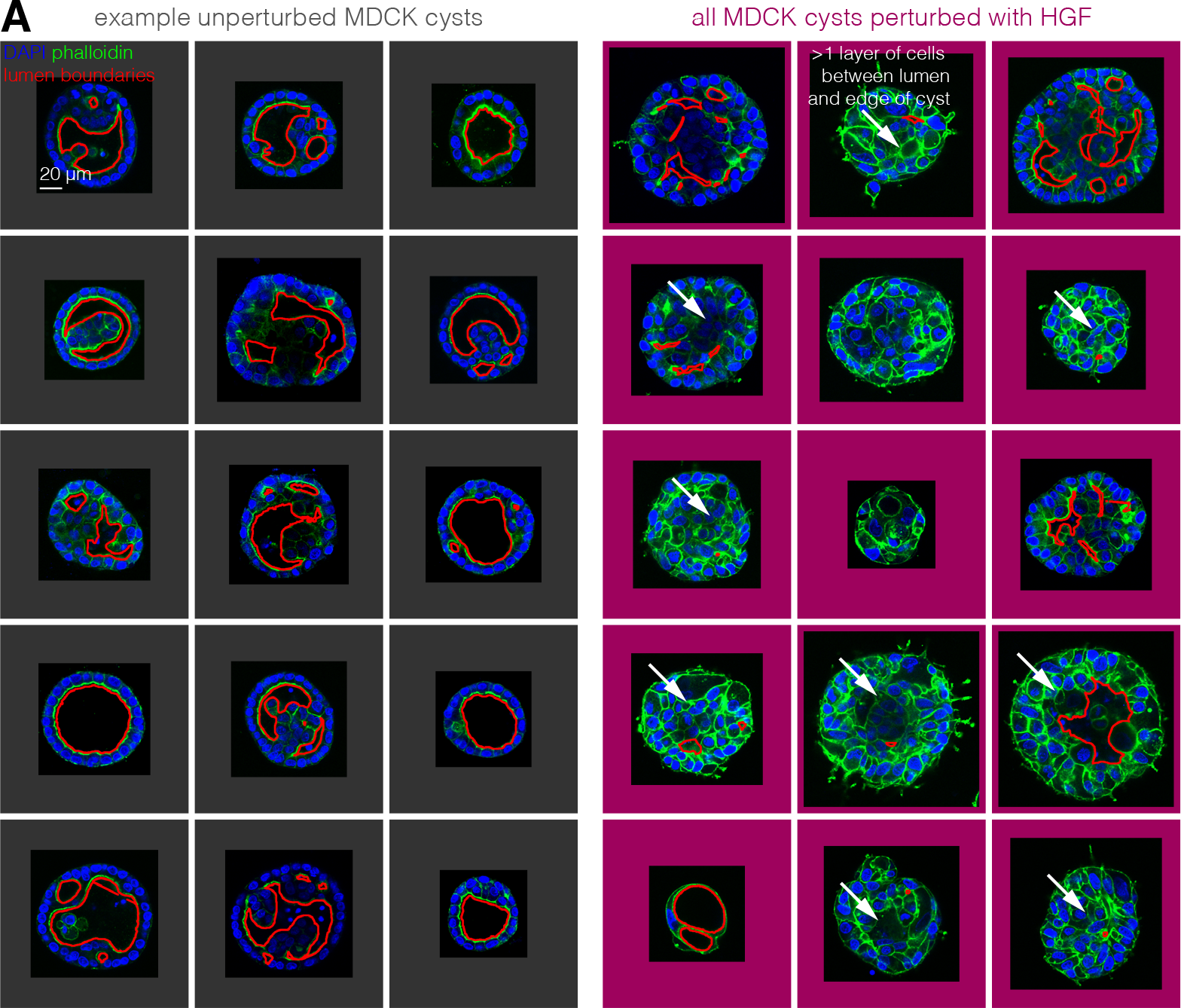
Spindle-like extensions are more common in MDCK cysts perturbed with HGF. **A.** 15 randomly-chosen unperturbed MOCK cysts and all 15 HGF-perturbed cysts. The boundary of the lumen is outlined in red. Spindle-like extensions are denoted with a white arrow.

**Supplemental Figure 21:**
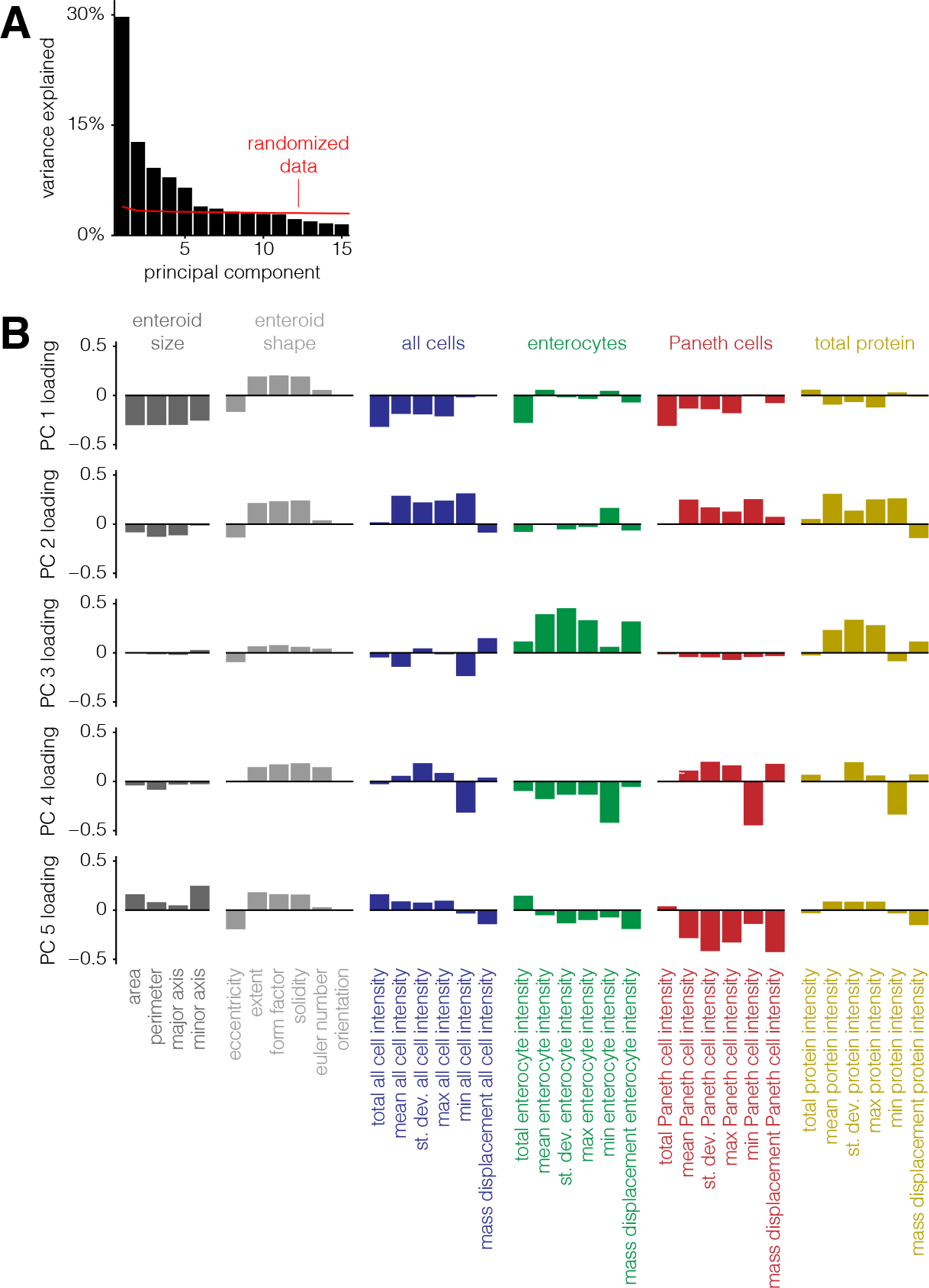
Enteroids obey a similar set of constraints to MDCK cysts. **A.** Variance explained by each principal component. The red line indicates how much variance is explained when the data is randomized before PCA (see methods for details). **B.** Loading of each feature on principal components one through five. Each feature is color-coded by what structure (enteroid size, enteroid shape, all cells, enterocytes, Paneth cells, or total protein) it describes.

